# Clade Distillation for Genome-wide Association Studies

**DOI:** 10.1101/2024.09.30.615852

**Authors:** Ryan Christ, Xinxin Wang, Louis J.M. Aslett, David Steinsaltz, Ira Hall

## Abstract

Testing inferred haplotype genealogies for association with phenotypes has been a longstanding goal in human genetics with several underlying challenges. A key advantage of these methods is the potential to detect association signals caused by allelic heterogeneity — when multiple causal variants modulate a phenotype — in both coding and noncoding regions. Recent scalable methods for inferring locus-specific genealogical trees along the genome, or representations thereof, have made substantial progress towards this goal; however, the problem of testing these trees for association with phenotypes has remained unsolved due to the growth in the number of clades with increasing sample size. To address this issue, we introduce several practical improvements to the kalis ancestry inference engine, including a general optimal checkpointing algorithm for decoding hidden Markov models, thereby enabling efficient genome-wide analyses. We then propose ‘LOCATER’, a powerful new procedure based on the recently proposed Stable Distillation framework, to test local tree representations for trait association. Although LOCATER is demonstrated here in conjunction with kalis, it may be used for testing output from any ancestry inference engine, regardless of whether such engines return discrete tree structures, relatedness matrices, or some combination of the two at each locus. Using simulated quantitative phenotypes, our results indicate that LOCATER achieves substantial power gains over traditional single marker testing and window-based testing in cases of allelic heterogeneity, while also improving causal region localization relative to single marker tests. These findings suggest that genealogy based association testing will be a fruitful approach for gene discovery, especially for signals driven by multiple ultra-rare variants.

**Author summary:** For a given set of individuals and at particular location in the genome, there is an underlying genealogical tree relating those individuals. Due to recombination, this tree is not static but rather varies along the genome. For decades investigators have sought to learn and use these trees to identify regions of the genome that impact human traits and disease. In other words, to find trait-associated trees where different clusters of relatives have, for example, high blood pressure. However, since these trees can be so enormous, it is difficult computationally to build them from DNA samples and difficult statistically to find trees with disease clusters: since each tree encodes so many possible clusters, it becomes hard to distinguish signal from noise. Here, we develop a new statistical method, LOCATER, to efficiently aggregate signals across disease clusters within each tree and thereby detect trait-associated trees. LOCATER can work with any ancestry inference method. We show LOCATER is better at detecting these trees than existing methods. We also introduce a suite of broadly applicable algorithms that make our ancestry inference software, kalis, and LOCATER computationally efficient. LOCATER is designed to work with any ancestry inference method.

## 1 Introduction

Recent heritability estimates predict that rare variants in regions with low linkage disequilibrium account for a substantial fraction of the unexplained (missing) heritability of common traits and diseases [1]. Since the statistical power to detect effects driven by rare variants is inherently limited by their low frequency, methods for identifying rare variant associations leverage allelic heterogeneity: the presence of multiple independent causal mutations affecting the trait of interest. These methods merge association signals from nearby rare variants under the premise that rare causal variants may be proximal to other causal variants [2, 3].

Mounting evidence suggests that allelic heterogeneity is quite common for human traits [4, 5]. Notably, a large-scale *in vitro* study following up on identified associations estimated that between 10% and 20% of expression quantitative trait loci (eQTLs) have multiple causal regulatory variants circulating in human populations [6]. Such results underscore the importance of these methods for defining new alleles and genes contributing to disease risk [7, 8, 1]. The opportunity for methods that can leverage allelic heterogeneity to identify overlooked associations will only increase with the size, population diversity, and sequencing depth of emerging genomic datasets.

Early groundbreaking association methods designed to harness allelic heterogeneity focused on testing inferred locus-specific genealogies, which provide a natural way of collecting independent association signals driven by nearby variants and imputing any unobserved variants [9, 10, 11]. While very elegant, these methods suffer from several technical limitations related to tree construction and statistical testing, such as requiring resampling or permutation to generate p-values, and ultimately have not been used much.

Recent advances in ancestry inference algorithms have made it possible to revisit genealogy-based trait association. Algorithms such as ARG-Needle [12], Relate [13], tsinfer [14], and kalis [15] have made it possible to perform local ancestry inference across the entire genome in modern datasets with hundreds of thousands or millions of samples. Given their accuracy in resolving recent genealogical relationships, inferred local ancestries are expected to be especially useful for detecting loci with multiple causal ultra-rare variants, which are signals that standard SMT will struggle to identify. This may be a particularly effective strategy for traits under strong purifying selection and cases where some of these ultra-rare causal variants correspond to complex hidden structural variations that are only observed in a given sequencing dataset via ultra-rare tagging variants that are far upstream or downstream. However, recent work aimed at applying these algorithms to improve disease mapping, most notably ARG-Needle [12], has focused on imputing hidden variants not explicitly observed in the original dataset and testing the inferred genotypes via single marker testing (SMT). Given the plummeting cost of high-coverage sequencing data and recent initiatives to improve structural variant detection and imputation [16], the number of missing variants is shrinking in modern datasets, limiting the gains available by using local ancestries to infer hidden variation.

Recent publications by Link et al. [17], Zhu et al. [18], and Gunnarsson et al. [19], are notable exceptions, showing a renewed interest in using genealogies to map genes by leveraging allelic heterogeneity. Building on earlier efforts like [9], these approaches target loci with allelic heterogeneity by using local ancestry inference methods to build a local genetic relatedness matrix for pre-specified windows along the genome or gene regions. These matrices are then tested for association with the phenotype of interest using a quadratic form test statistic. This statistical approach mirrors SKAT and more recent methods in rare-variant gene-based testing that also aim to harness allelic heterogeneity to gain statistical power [2, 20]. However, due to the inherent sensitivity of quadratic form test statistics to the presence of many non-causal (null) variants [21], these approaches struggle to maintain statistical power under the enormous multiple testing burden incurred when testing all of the clades in a local tree for trait association.

Existing rare variant association methods that aim to leverage allelic heterogeneity by testing collections of variants, such as STAAR, limit their multiple testing burden by using functional information, such as gene coding sequences, to define restricted sets of variants or more flexibly down-weight certain variants [2, 20]. This approach has been applied to many different sequencing based studies, and has proven to be a fruitful approach for identifying new gene-phenotype associations across a multitude of traits [22]. Despite their success in coding regions, it has proven difficult to extend rare variant association tests beyond coding regions where the majority of biologically critical signals are found. 90% of GWAS hits for common diseases lie in non-coding regions, at a median distance of 36 kb from the nearest TSS. [23, 24]. It is unclear how one should define collections of variants in non-coding regions; sliding windows are the standard approach [20].

Outside of a gene’s coding region, which in humans has a median length ≈ 3Kb, there is a much larger regulatory region over which causal variants may be dispersed, complicating the use of sliding windows. Ideally, one would try to incorporate many variants over a genomic region in order to maximize the chance of aggregating signals from more than one causal variant. However, including too many non-causal (null) variants diminishes statistical power, and variant impact prediction, which could be used to narrow down variants to test in a given sliding window, remains extremely challenging in noncoding regions.

There are “sparse-signal” statistical methods, often deployed alongside quadratic forms, that aim to improve power in the presence of many null variants. Notable examples include the ACAT routine built into STAAR [20] and Generalized Higher Criticism [25]. However, these sparse-signal methods do not distinguish between a variant set where two highly-linked variants are observed to be associated with the phenotype and a variant set where two unlinked variants are observed to be associated with the phenotype. In the highly-linked case, one variant is essentially a proxy for the other and we have only one association signal. In the unlinked case, the signals coming from the two independent variants serve as independent pieces of evidence against the null hypothesis and should be combined. In order to control the type-I error in the highly-linked case, ACAT cannot combine signals across variants with high efficiency, which yields a loss of power in the unlinked case. This simple two-variant argument extends to the case where we may be attempting to combine association signals across several variants.

We recently proposed a general statistical approach, Stable Distillation (SD), which can distinguish between the highly-linked and unlinked case [21]. There, in a gene-testing example using simulated data, we used SD to explicitly model the dependence structure between variants and achieved increased power over ACAT and related methods as a result. Building on SD, here we present a general framework, LOCATER, for trait association based on inferred local genealogies in both coding and non-coding regions. Here we focus exclusively on testing quantitative traits, although we plan to extend LOCATER to binary traits in future work.

LOCATER is designed to work in conjunction with any ancestry inference engine of the user’s choosing with an easy-to-use API available through our `LOCATER` package for the `R` language [26]. Modern ancestry inference methods typically represent local ancestries as discrete trees (perhaps with probabilistic weights on the edges), local relatedness matrices, or some combination of the two. Examples of discrete tree inference methods include `tsinfer`. These clades may have probabilistic weights, as provided by recent probabilistic ARG inference methods such as SINGER [27]. On the other hand, ancestry may also be represented in terms of local pairwise relatedness, typically summarized as a local relatedness matrix, as produced by `kalis, Relate`, and *Gamma-SMC* [28]. A set of observed haplotypes can typically be explained by an enormous number of underlying tree topologies, especially once we look beyond the recent past; pairwise methods account naturally for this topological uncertainty. As described in Section 2.1, LOCATER provides a framework for boosting SMT results with independent association signals based on local ancestry represented in either, or both, of these two forms produced by any ancestry inference engine. In order to highlight this feature, in this paper we apply LOCATER to complimentary discrete clade and matrix-based representations of local ancestry obtained via the local ancestry inference engine `kalis`. Although `kalis` does not scale as well as alternatives like `tsinfer`, a probabilistic model allows us to limit statistical testing to clades that have substantial evidence of existing at a locus of interest, thereby conserving statistical power. SINGER may provide a strong probabilistic alternative in future studies. The algorithmic improvements to `kalis` that we present in this paper, including an optimal checkpointing routine for discrete-time hidden Markov models (HMMs) and linear-time clustering algorithm, may be useful for accelerating alternative models.

Our focus on using genealogies to boost SMT signals rather than testing gene windows or sliding windows along the genome is another key point of departure of LOCATER from existing work. While our approach may appear ill-advised at first, given that it requires LOCATER to clear the standard SMT Bonferroni threshold (≈ 10^−8.5^), it leverages the statistical efficiency of SMT against sparse signals. This approach also removes questions of window size and step length, which is a natural advantage of using genealogies in the first place. As we will demonstrate below, by returning “genealogy-boosted” SMT signals, LOCATER also aids the localization of causal variants.

## 2 Results

LOCATER assumes that genome-wide SMT has already been performed. This is done simply to avoid the computational burden of inferring ancestries and running LOCATER at every locus. We focus on the subset of variants with putatively significant SMT results (e.g., *P*< 10^−4^) and compute the local ancestry at each of those variants. At each target variant, LOCATER then takes the residuals from the SMT and tests any inferred discrete clade structure with Stable Distillation (SD) [21]. SD naturally returns a new set of residuals which are guaranteed to be independent of the original SMT p-value and the p-value returned by SD under the null hypothesis. We then pass this set of residuals to any quadratic-form based method that tests the pairwise-relatedness structure inferred at the variant of interest. The resulting three independent p-values may then be combined to obtain a potentially boosted signal at a locus with allelic heterogeneity. This approach makes it straightforward to integrate LOCATER into the analysis of genome-wide association results. Of course, the resulting p-values must still be compared against a genome-wide multiple testing threshold as if LOCATER was run at every candidate variant. LOCATER can easily be applied in special cases where the inferred ancestry at a locus only comes in the form of discrete clades (eg: a tree) or pairwise-relatedness (eg: a local relatedness matrix). In both our SD procedure and quadratic form testing procedure, we have developed scalable methods to adjust for population structure and background covariates (see Methods).

Below we introduce the LOCATER model. We then proceed to describe our methodological contributions in three parts: generating the ancestry representations required by LOCATER using the new routines we have introduced in an update to `kalis` [15], making LOCATER fast and robust to population structure, and efficiently combining the p-values returned by LOCATER. Finally, we demonstrate the calibration and power of LOCATER via simulation.

### 2.1 The LOCATER Model

Consider a genomic dataset with *n* participants phased in segments along the genome, each segment consisting of *N* = 2*n* phased haplotypes along an entire or subsection of a chromosome with a total of *V* variants. Although we only address the diploid case in this paper, our approach may be readily extended to non-diploid organisms. Below we will consider testing each variant within a given segment for association with some quantitative phenotype of interest *Y* ∈ ℝ^*n*^. When determining genome-wide significance thresholds, the total number of candidate variants across all genomic segments must be accounted for. However, in order to conserve computational resources, as depicted in Figure 1, we may only be interested in a subset of candidate variants within each segment based on preliminary SMT results or other genomic annotations. We call this set of target loci ℒ ⊆ [*V*] and index them by their position along a given segment sequentially from *ℓ* = 1, …, *L* = |ℒ|.

**Figure 1:**
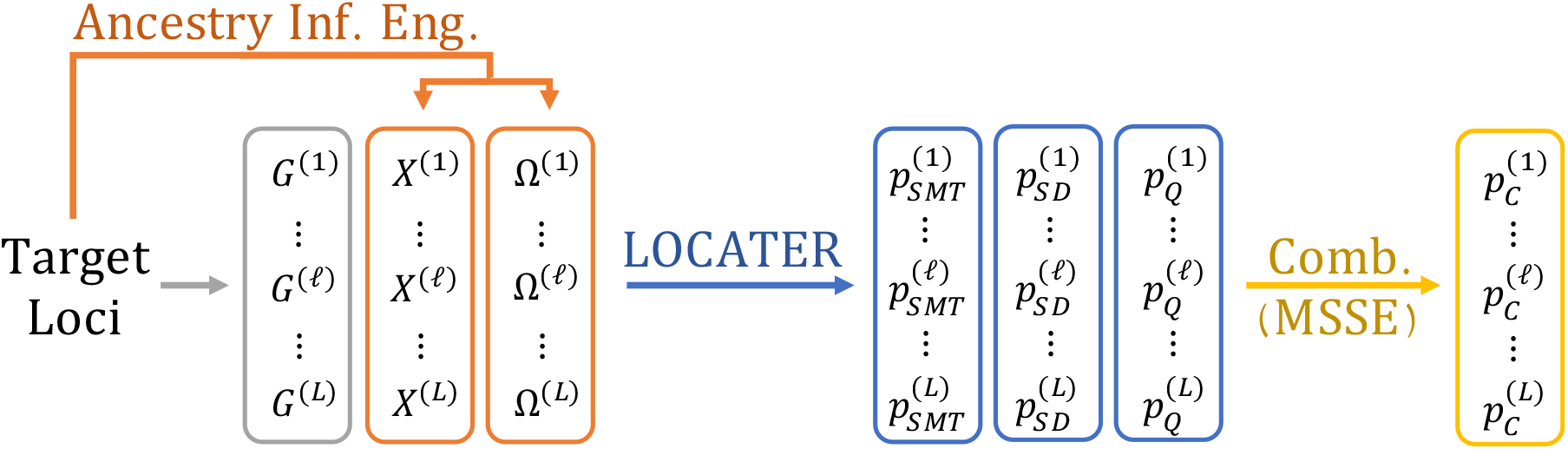
The LOCATER Pipeline. We begin ancestry-based association testing with a set of putatively interesting target loci, typically identified via single marker testing, indexed {1, …, *L*}. At each target locus *ℓ*, we extract the genotype vector *G*^(*ℓ*)^ ∈ {0, 1, 2}^*n*^ and use an ancestry inference engine to infer local clade genotypes *X*^(*ℓ*)^ ∈ {0, 1, 2}^*n*×*p*^ and/or a local relatedness matrix Ω^(*l*)^ ∈ ℝ^*n*×*n*^. We then use LOCATER to calculate three p-values testing whether *G*^(*ℓ*)^, *X*^(*ℓ*)^, or Ω^(*l*)^ predict the phenotype respectively. These three p-values are guaranteed to be independent under the null hypothesis, so they may be easily combined with many methods, in this paper we propose and use MSSE 4.8, to obtain a combined ancestry-association p-value 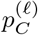 at each target locus.

Let *A* ∈ ℝ^*n*×*q*^ be a matrix of background covariates and *G*^(*ℓ*)^ ∈ {0, 1, 2}^*n*^ be the genotype vector observed at locus *ℓ*. Depending on the type of inference engine used to infer the local ancestry structure at locus *ℓ*, we may have a set of inferred genotypes *X*^(*ℓ*)^ ∈ {0, 1, 2}^*n*×*p*^ corresponding to edges in a tree inferred at *ℓ*, a local relatedness matrix Ω^(*l*)^ ∈ ℝ^*n*×*n*^ inferred at *ℓ*, or both. Here we tackle the general case assuming that our ancestry inference engine has returned both *X*^(*ℓ*)^ and Ω^(*ℓ*)^, each capturing different parts of the ancestral structure at locus *ℓ*. Our approach can easily be applied to the special cases where only *X*^(*ℓ*)^ or Ω^(*ℓ*)^ are available.

For a set of parameters *α* ∈ ℝ^*q*^, *γ* ∈ ℝ, *β* ∈ ℝ^*p*^, and *τ* ∈ ℝ_≥0_, LOCATER assumes the following model for a quantitative phenotype vector *Y* ∈ R^*n*^.

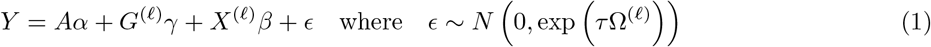

Here, exp denotes the matrix exponential. Under this model we test whether genetic variation at locus *ℓ* affects phenotype *Y* by testing the null hypothesis 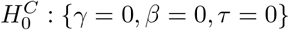. Here and for the rest of the paper, we will assume that *Y* has been obtained using the rank-matching procedure described in Section 6.6. This normalization ensures that the residuals of *Y* have unit variance under 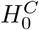, justifying the absence of a variance scale parameter (typically denoted as *σ*^2^) in Equation (1).

LOCATER tests 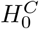 by decomposing it into three sub-hypotheses. First, we use the standard SMT to test 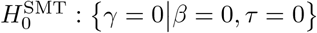, yielding a p-value *p*_SMT_. Then we test whether any of the locally inferred clades predict the phenotype 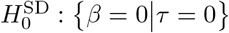 using SD, yielding a p-value *p*_SD_. Finally, we test whether any remaining local ancestry structure encoded in Ω^(*ℓ*)^ affects the phenotype by testing 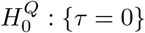 with a quadratic form test statistic, yielding a p-value *p*_Q_. We further describe the explicit routines LOCATER uses to test 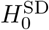 and 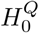 in Section 2.3. When these three sub-hypotheses are tested in this order, the independence guarantees of SD ensure that the resulting p-values (*p*_SMT_, *p*_SD_, *p*_Q_) are mutually independent under the null hypothesis 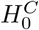. Thus, after running LOCATER, the user may combine these three p-values using any valid method for aggregating independent p-values. Here we propose a variant of Fisher’s method that we call Maximizing over Subsets of Summed Exponentials (MSSE) which yields more power in this setting (see Section 4.8). Figure 1 overviews the role of LOCATER in the context of an ancestry-based association testing pipeline. Next we delineate how the algorithms we have implemented in `kalis` v2 allow us to rapidly obtain *X*^(*ℓ*)^ and Ω^(*ℓ*)^ across target loci in our present study.

### 2.2 Algorithmic Advances in kalis v2

#### 2.2.1 Local Genealogy Inference with kalis

`kalis` [15] provides a high-performance implementation of various versions of the Li & Stephens (LS) haplotype copying model which have become ubiquitous in modern genomic analysis [29, 30]. Here we overview our novel algorithmic contributions to a new release, `kalis v2`, which together allow us to efficiently calculate *X*^(*ℓ*)^ and Ω^(*ℓ*)^ sequentially at a given set of target loci *ℓ* = 1, …, *L* so that they can be tested downstream using LOCATER. In order to explain these contributions, we begin with a brief overview of local ancestry inference using `kalis`.

As with all ancestry inference engines, the ancestry at a given target locus *ℓ* is learned based on the observed genomic variation upstream and downstream of *ℓ*. Since the LS model is a special case of an HMM, ancestry information provided by variants upstream of *ℓ* can be summarized by the forward probabilities at *ℓ*; and variants downstream by the backward probabilities at *ℓ* [31]. Given a set of *N* haplotypes and a single target locus *ℓ*, `kalis` implements the forward algorithm to iterate over variants upstream of *ℓ*, starting at the left end of the genomic segment, to obtain a matrix of forward probabilities *f*^(*ℓ*)^ ∈ ℝ^*N*×*N*^ at *ℓ*. Similarly, `kalis` implements the backward algorithm to iterate over variants downstream of *ℓ*, starting at the right end of the genomic segment, to obtain a matrix of backward probabilities *b*^(*ℓ*)^ ∈ R^*N*×*N*^ at *ℓ*. Each column 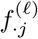 and column 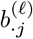 corresponds to an independent LS HMM where we model recipient haplotype *j* as a mosaic of the other *N* − 1 haplotypes in the sample. This independence allows `kalis` to compute the columns of *f*^(*ℓ*)^ and *b*^(*ℓ*)^ in parallel and exploit modern compute architectures. See [15] for further details. The product 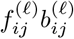 can be interpreted as proportional to the probability that recipient haplotype *j* “copies” from donor haplotype *i* at locus *ℓ* under the LS model. By definition, 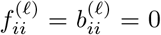 for all haplotypes *i*. `kalis` makes *f*^(*ℓ*)^ and *b*^(*ℓ*)^ easily and rapidly accessible in the R language [26] for downstream computation, with all time-critical code written in high performance C.

Along the lines of [13] and [15], we define the distance from haplotype *j* to haplotype *i* as

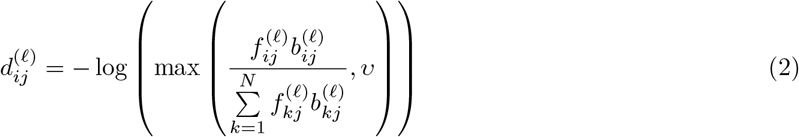

where *υ* ≈ 4.94 × 10^−324^ to guard against underflow to zero with double precision floating point arithmetic. For efficiency the distance matrix 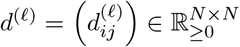 is never explicitly constructed, but it is implicitly used to construct *X*^(*ℓ*)^ and Ω^(*ℓ*)^ for testing with LOCATER, as further delineated below.

Throughout this paper, we use `kalis` to run the modified LS model used in Relate [13]. See Section 6.1 for further details. This modified model leverages ancestral allele information to improve local genealogy inference. In this paper we only simulate phased genomic datasets where the ancestral allele of each variant is known; this is a feature of our chosen ancestry inference engine and not a general requirement of LOCATER. Under this modified LS model, Speidel et al. showed that the distance 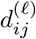 will be proportional to the number of proximal variants that differ between haplotype *i* and haplotype *j* in non-recombining segments and that the full matrix *d*^(*ℓ*)^ yields consistent local ancestry inference.

#### 2.2.2 Optimal Checkpointing

Especially when processing many phenotypes in parallel, the number of target variants along a given genomic segment, *L*, may be very large. Since the amount of memory required to store a local relatedness matrix Ω^(*ℓ*)^ at a given target locus scales 𝒪(*n*^2^), storing these matrices at any appreciable number of variants quickly becomes untenable: even in the case where *n* = 30, 000 samples (a scale we will consider in our simulations), 28.8 GB of memory is required to store a single Ω^(*ℓ*)^. In particular, offloading from memory may be impractical as there would be a considerable time cost to writing and reading Ω^(*ℓ*)^ to and from disk. To avoid storing any Ω^(*ℓ*)^, we will take a “test-it-and-forget-it” approach: we will obtain *X*^(*ℓ*)^ and Ω^(*ℓ*)^ at one target variant at a time and test both *X*^(*ℓ*)^ and Ω^(*ℓ*)^ with LOCATER before moving on to the next target variant in ℒ. This “test-it-and-forget-it” approach is only computationally tractable due to a new, general checkpointing algorithm for discrete-time HMMs that we have introduced in `kalis v2`. In short, our checkpointing approach allows us to infer local relatedness matrices sequentially across consecutive target loci while minimizing the computation required subject to a fixed memory budget.

To understand the need for checkpointing, as mentioned above and further detailed below, recall that we implicitly need the pairwise distance matrix *d*^(*ℓ*)^ from Equation (2) returned by `kalis` in order to obtain *X*^(*ℓ*)^ and Ω^(*ℓ*)^ at each target locus *ℓ*. While `kalis` v1 can efficiently propagate the forward and backward recursions to obtain the forward probability matrix *f*^(*ℓ*)^ and backward probability matrix *b*^(*ℓ*)^ needed to calculate *d*^(*ℓ*)^ at a single locus *ℓ*, obtaining the pair *f*^(*ℓ*)^, *b*^(*ℓ*)^ at sequential positions *ℓ* is challenging for HMMs due to the uni-directionality of the forward and backward recursions. The compute-minimizing approach would involve running a single pass of the forward algorithm – iterating the forward recursion to target locus 1, then locus 2, and so on until locus *L* – and a single pass of the backward recursion from target locus *L* to target locus 1 while storing *f*^(*ℓ*)^ and *b*^(*ℓ*)^ at every *ℓ* = 1, …, *L*. Since each *f*^(*ℓ*)^ and *b*^(*ℓ*)^ consumes 8*N*^2^ bytes of memory (e.g.: 80 GB for *n* = *N/*2 = 50, 000 haplotypes), this approach requires far too much storage for most genomic datasets. On the other hand, we have the memory-minimizing approach, where we restart the forward and backward recursions from the respective ends of the genomic segment for every target locus. While this approach only requires storing a single *f*^(*ℓ*)^ and a single *b*^(*ℓ*)^ at any given time, it demands far too much compute time for most genomic datasets, requiring 𝒪(*L*^2^*N*^2^) floating point operations (FLOPs) — a prohibitive cost. An attempt to rescue this approach by splitting the genome into smaller segments (running in chunks) would still require 𝒪(*L*^2^*N*^2^) compute time.

We provide a checkpointing algorithm that finds an optimal balance in this memory-compute trade-off, minimizing the compute time required given a fixed memory budget. The overall idea is to occasionally stop the forward recursion and store *f*^(*ℓ*)^ at its current position as a checkpoint (typically overwriting an old checkpoint) in order to avoid repeatedly restarting the forward recursion from the beginning of the genomic segment. Here we provide a broad overview of our checkpointing approach. We start with a user-specified memory budget capable of holding *C* checkpoints, each storing a *N* × *N* matrix of forward probabilities *f*^(*ℓ*)^. We run the backward recursion once across the entire chromosome or genomic segment, stopping at each consecutive target locus sequentially from the target locus with the largest position (*ℓ* = *L*) to the target locus with the smallest position (*ℓ* = 1). When the backward recursion stops at a given target locus, we run the forward recursion from the nearest checkpoint to meet the backward recursion and so obtain *X*^(*ℓ*)^ and Ω^(*ℓ*)^ at that target locus. Note it is natural for us to perform this backwards along the genomic segment, since there is a slightly higher computational cost for the backward recursion and hence we favor repetitive restarts of the forward recursion.

Iterating from locus *L* down to locus 1 makes minimizing the compute required for the backward recursion trivial: we simply visit each locus sequentially in a single pass. The challenge is determining where and when to overwrite existing checkpoints to minimize the total distance (number of variants) that the forward algorithm needs to iterate over in order to provide forward matrices in reverse order *f*^(*L*)^ → *f*^(*L*−1)^ → · · ·. In Section 4.3 we show how to solve for a schedule of checkpoints that achieves this minimum for any discrete time HMM, given storage for a fixed number of checkpoints *C*. We call this solution the “optimal checkpointing schedule.” After a forward matrix *f*^(*ℓ*)^ is obtained at a given target locus *ℓ*, this schedule instructs `kalis` which checkpoint to use to restart the forward recursion to obtain the next forward matrix at locus *ℓ* − 1, and where to lay down new checkpoints (if any) as the forward recursion proceeds to that locus. The checkpointing schedule also dictates where to initialize the *C* checkpoints as we iterate the forward recursion to the first target locus *L*. As shown in Supplementary Figure 5, the use of only 5 to 10 checkpoints already reduces the required number of FLOPs orders of magnitude closer to the best possible complexity, which would be 𝒪(*N*^2^*L)* given substantial memory.

As detailed in Section 4.3, solving for the optimal checkpointing schedule can be computationally intensive for any given set of target loci. The version of the checkpointing schedule solver currently implemented in LOCATER assumes that target loci are evenly spaced. This simplification does not qualitatively change performance but allows us to solve for the optimal checkpointing strategy for a given *L* via a dynamic program, making the solution readily available. Our checkpointing implementation is available via the `ForwardIterator` function and associated helper functions now provided in `kalis v2`.

Of course, for datasets with a large number of samples, there may not be sufficient capacity to store many checkpoints in memory. At a minimum, running `kalis` on *n* samples (2*n* phased haplotypes) to obtain each Ω^(*ℓ*)^ requires 32*n*^2^ bytes to store the forward and backward probabilities and another 8*n*^2^ bytes to store Ω^(*ℓ*)^. Storing each additional checkpoint of forward probabilities requires 16*n*^2^ bytes. Given the nested nature of our checkpointing algorithm, most checkpoints can be stored on disk rather than memory, which comes at minimal computational cost as long as one or two of the checkpoints (the ones that are closest to the current target loci) are always kept in memory. We plan to add native support for storing file-backed checkpoints to `kalis` in the near future. Looking further ahead, `kalis` can already be distributed across machines, each running the LS model on a different subset of recipient haplotypes [15], but running LOCATER across distributed machines would require substantial network communication. Reducing this communication is a direction of future work.

#### 2.2.3 Calculating Inferred Clade Genotypes from the LS Model

The LOCATER model (Equation (1)) admits a matrix of genotypes *X*^(*ℓ*)^ encoding any clades (marginal tree edges) inferred at locus *ℓ*. In principle, given the distance matrix *d*^(*ℓ*)^ obtained via `kalis` at locus *ℓ* (Equation (2)), any number of clustering algorithms could be used to infer a marginal tree topology from *d*^(*ℓ*)^. For example, `Relate` clusters a normalized version of *d*^(*ℓ*)^ with average linkage (UPGMA) [13]. From the resulting tree topology, one could then encode each of the inferred clades (or some subset of them) via *X*^(*ℓ*)^. While this is a promising approach for future work, in this paper, we only stored clade genotypes corresponding to very small inferred clades (each typically including 2 to 10 haplotypes) in *X*^(*ℓ*)^ and encode all larger-scale relatedness structure in Ω^(*ℓ*)^. Focusing on just these rare clades, which we will refer to as “sprigs,” rather than all of the clades in the tree, allows us to showcase the flexibility of LOCATER — the ability of LOCATER to incorporate hard-called clades via *X*^(*ℓ*)^ and remaining relatedness structure via Ω^(*ℓ*)^.

We identify sprigs based on the neighborhood — i.e., the set of tied nearest-neighbors — of each haplotype *j*:

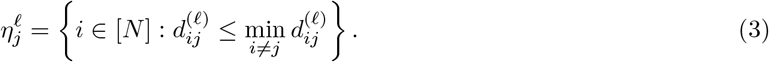

Note that, by this definition, haplotype *j* is always a nearest neighbor of itself. In practice we obtain these neighborhoods as a by-product of the clustering algorithm we use to construct Ω^(*ℓ*)^ (see Section 2.2.4). We implicitly use the collection of neighborhoods 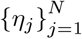 to construct an undirected graph where each haplotype is a vertex, and edges connect haplotypes that agree on being in each other’s neighborhood. We use a greedy clique-finding procedure over the nearest neighborhoods to rapidly identify maximal cliques within this implicit graph. Haplotypes within each clique are assumed to belong to the same sprig, yielding sprig genotypes that we encode in *X*^(*ℓ*)^.

#### 2.2.4 Calculating Relatedness Matrices from the LS Model

Having encoded the locally inferred sprig genotypes in *X*^(*ℓ*)^, we summarize all of the remaining genealogical structure in *d*^(*ℓ*)^ via Ω^(*ℓ*)^. While LOCATER can accept any real symmetric matrix Ω^(*ℓ*)^, in order to optimize power we model our choice of Ω^(*ℓ*)^ in this paper after the expected genetic relatedness matrix (eGRM) as presented in Equation 1 of [17]. We cannot directly use the definition of the eGRM given in [17] because the construction there requires a set of discrete clade calls. Constructing a local relatedness matrix from the distances *d*^(*ℓ*)^ is more complicated because, as described in Section 2.2.1, each column is calculated using an independent LS HMM. Thus, different columns of *d*^(*ℓ*)^ may disagree on the exact boundaries of particular clades in the underlying genealogy. This is a general feature of ancestry inference methods that work in a parallel or pairwise fashion across haplotypes. Rather than overriding the LS model and using hierarchical clustering or some other approach to try to align these clade calls, here we generalize the eGRM to allow for this asynchrony. Our generalization, presented in Section 4.4, expresses an eGRM in terms of asymmetric distances like those provided by *d*^(*ℓ*)^ while naturally allowing for such unaligned probabilistic clade calls.

This generalization of the eGRM requires us to use the distances within each column of *d*^(*ℓ*)^ to call a set of nested neighborhoods around the corresponding haplotype. Calling these nested neighborhoods amounts to clustering the distances in each column of *d*^(*ℓ*)^. In order to do this efficiently for large *n* datasets, we developed a general, multithreaded, single-pass algorithm based on doubly linked lists to cluster real numbers on a closed interval when clusters must be separated by some fixed minimum distance. This approach allows us to cluster each column of *d*^(*ℓ*)^ in 𝒪(*N*) time. In experiments on simulated haplotype data, we achieve roughly an order of magnitude speedup over merge sort. In order to conserve memory, our implementation does not explicitly store the clustering results for each column of *d*^(*ℓ*)^. Rather, we use these clusters to directly construct columns of an asymmetric version of Ω^(*ℓ*)^ on the fly, directly collapsing haplotype level relatedness down to sample level relatedness as we go. Taking the symmetric part of the resulting matrix gives us Ω^(*ℓ*)^. See Section 4.5 for details and the specific construction of Ω^(*ℓ*)^ we use in this paper.

As a by-product of the clustering used to construct Ω^(*ℓ*)^, we also return the nearest-neighbor set of each haplotype, which is then used to call sprigs and construct *X*^(*ℓ*)^ (see Section 2.2.3). After calling sprigs using these neighborhoods, in order to avoid testing these sprigs in both *X*^(*ℓ*)^ and Ω^(*ℓ*)^, we efficiently remove the structure associated with those sprigs from Ω^(*ℓ*)^ using some additional statistics reported by our clustering algorithm before passing *X*^(*ℓ*)^ and Ω^(*ℓ*)^ on to LOCATER for testing. All of these methods are available in `kalis v2`.

### 2.3 LOCATER Testing Routines

All of LOCATER’s routines have been written in terms of matrix operations, allowing multiple quantitative traits to be tested in parallel with minimal additional computational cost. This includes the first implementation of a parallelized SD algorithm. For a given phenotype, this SD algorithm yields decoupled estimators for the effect *β*_*j*_ of each inferred clade genotype 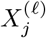. We then combine the independent two-sided p-values corresponding to these independent estimators via the Rényi Outlier Test [32] to obtain 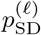 at each locus. As demonstrated in [21], this approach yields considerable gains in power over alternative methods when very few (but more than one) of the *β*_*j*_ are non-zero; in other words, when more than one of the inferred genotype clades is associated with the phenotype. See Section 4.6 for details about the specific SD procedure we use.

LOCATER also deploys several statistical and algorithmic innovations to calculate 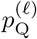. Under the LOCATER model (Equation (1)), the score statistic against the null *τ* = 0 is a quadratic form,

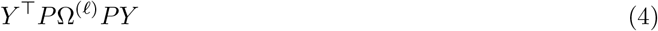

where *P* = *I* −*QQ*^⊤^ and (*A, G*^(*ℓ*)^) = *QR* is the QR decomposition adjusting for the background covariates and the tested genotype *G*^(*ℓ*)^. In order to avoid launching unnecessary and expensive partial eigendecomposition routines at every target variant, we use a series of approximations to first assess whether the combined LOCATER p-value 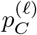 is sufficiently small to be interesting across any of the phenotypes. When it is, further eigendecomposition of *P*Ω^(*ℓ*)^*P* is deployed in order to obtain precise estimates.

We found that the Satterthwaite approximation [33], which is commonly used for testing quadratic forms [34], did not yield robust tail probability estimates for LOCATER. This may be because in this setting the matrices *P*Ω^(*ℓ*)^*P* are typically close to, but not quite, positive semi-definite — a key assumption of the Satterthwaite approximation. We overcame this obstacle with a new, robust tail approximation method for quadratic forms based on a shifted difference of chi-square random variables. In combination with our approximation stopping criteria, this tail approximation provides a basis for emerging genealogy-based association methods to reliably test local pairwise relatedness matrices, which may often not be positive semi-definite. Our tail approximation method has the added advantage that it naturally admits three parameters — 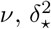, and 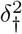 — to help control for population structure (in a way that generalizes genomic control to quadratic forms). Importantly, these three parameters were chosen to be orthogonal to the spectral parameters governing the distribution of Equation (4). If any inflation is observed in the Q-Q plot of 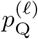 p-values after running LOCATER, this orthogonal parameterization allows us to adjust 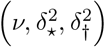 and rapidly calculate new 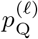 p-values without requiring the re-estimation of local ancestries at any target locus. Please see Section 4.7 for further details.

We also use a novel multi-threaded algorithm for efficiently projecting out background covariates when calculating the matrix traces needed for these tail approximations, which is further described in Section 6.4. All final p-values *p*_Q_ involving eigenvalue terms are calculated using the FFT-based approach implemented in the R package `QForm` [35].

Finally, we combine our three p-values, 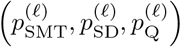, using a modified version of Fisher’s combination test we call MSSE, as described in Section 4.8.

### 2.4 Type-I Error Control

In order to confirm the calibration of LOCATER empirically, we simulated 1k independent genomic datasets, each consisting of 30k samples of a 1Mb chromosome. See Section 4.1 for more details. For each dataset, we simulated 1k independent phenotype vectors assuming no causal variants. See 4.2 for more details of our phenotype simulation approach. This yielded a total of 1 million independent phenotype vectors. To simulate each phenotype, we first sampled two background covariate vectors: *a*_1_ from independent standard Gaussian random variables and *a*_2_ from independent Rademacher random variables. We then sampled the phenotype vector from *Y* ∼ *N* (*a*_1_ + *a*_2_, *I*). We tested each phenotype vector for association with the ancestry inferred at the mid-point of the corresponding chromosome using LOCATER. We display a Q-Q plot for − log_10_ of those p-values, as well as for each LOCATER sub-test — SMT, SD, and QForm — in Section 5.2 (Supplementary Figures 6,7,8,9). These Q-Q plots confirm that the p-values returned by each sub-test and the combined LOCATER p-value are all well calibrated under the null hypothesis.

### 2.5 Power

We compared LOCATER to standard SMT across a variety of genetic architectures. We assessed every possible combination of the following causal variant assumptions. We considered 3, 9, or 15 causal variants based on the number of independent causal alleles that were observed in the large follow-up study of GWAS hits by Abell et al. (see Figure 4b of [6]). We also considered causal variants with any derived allele count, derived allele count of 2 (doubletons only), or intermediate variants with derived allele count in [150,750). That is equivalent to a derived allele frequency (DAF) in [0.0025,0.0125). Lastly, we considered the case where all causal variants are observed or all causal variants are hidden. This yielded a total of 18 genetic architectures. Note that by ‘observed’, we mean that the causal variants were included in the dataset passed to each association method; by ‘hidden’, that they were not included in the dataset passed to each association method and thus could only be inferred via LD. In each simulation, causal variants were randomly assigned from among those fulfilling the required allele count requirements within a 10 kb window in the center of each simulated 1 Mb segment.

Under each genetic architecture, we estimated power as a function of the underlying total association signal strength: the − log_10_ p-value that one would obtain by testing the simulated phenotype *Y* with an oracle ANOVA model that “knows” which are the active predictors and targets only those for testing. To improve the interpretability of our power curves, following [21], we used the *QR*-decomposition to ensure that the total association signal was evenly split among the causal variants in every simulation. In other words, we ensured that the *observed* contribution of each causal variant to the total association signal was essentially equal for each simulated *Y*. Please see Section 4.2 for more details of our phenotype simulation approach.

In calculating power, we count our causal region as “discovered” if a testing method has a p-value less than the 10^−8.5^ threshold *anywhere* along the entire 1Mb region. This definition reflects how new associations are discovered in practice and provides a relatively strict benchmark. Each point of the resulting power curves was estimated via 1k independent samples: we simulated 10 independent phenotype vectors for each of 100 independent genomic datasets, each consisting of a 1 Mb chromosome sampled for 30k individuals. These power curves are available in Section 5.3 (Supplementary Figures 10,16,22,13,19,25). We summarize each of these curves with the estimated minimum signal strength required to achieve 80% power (lower is better). Figure 2 displays those estimates for LOCATER and SMT across all simulations where the underlying causal variants were observed; Figure 3, for those hidden.

**Figure 2:**
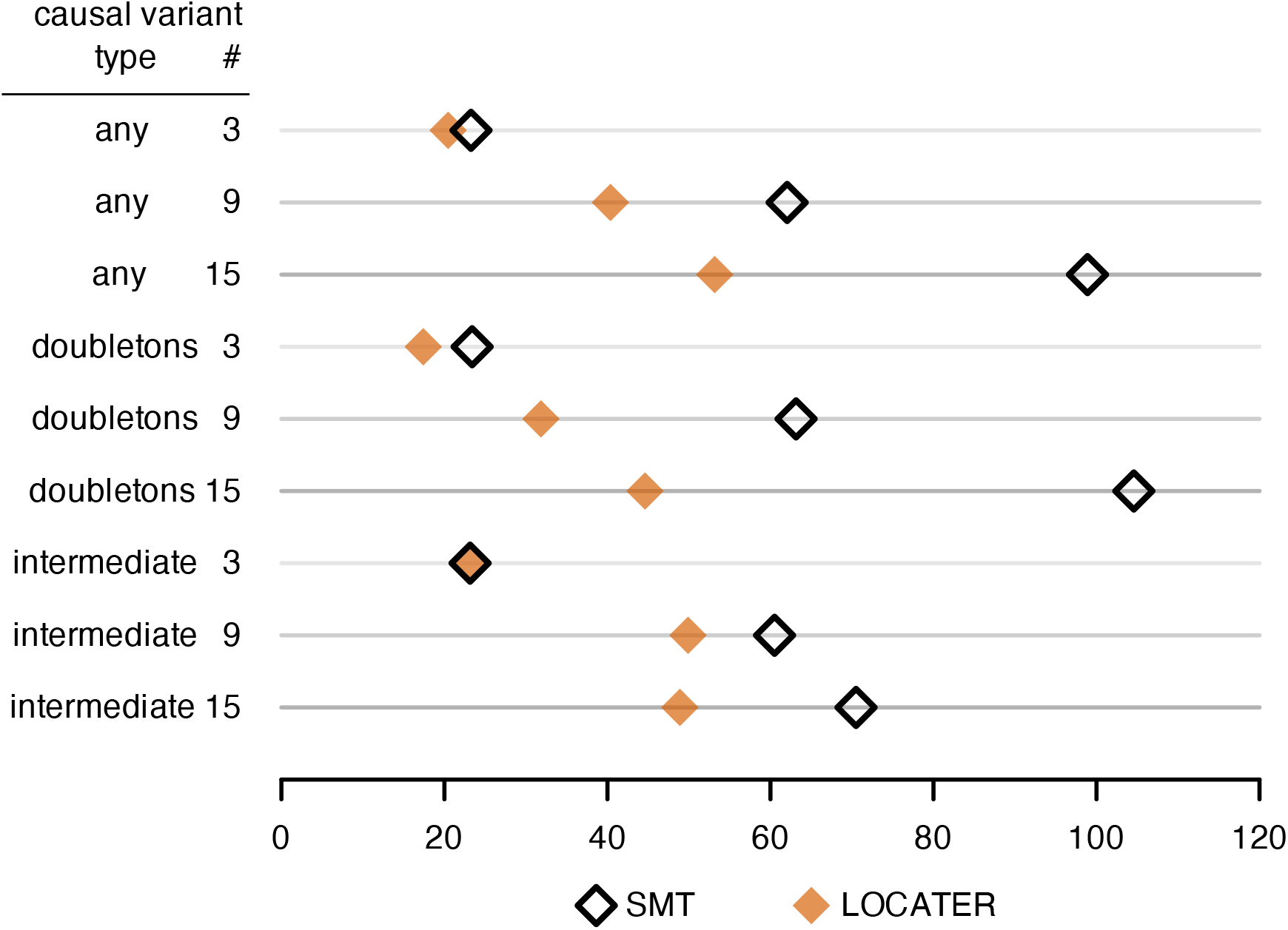
Dotplot of total association signal strength required to achieve 80% power (lower is better) under various simulation conditions where all causal variants were observed. Causal variant # denotes the number of simulated causal variants. Causal variant type “any” means any variant could be causal; “doubletons” means only doubletons could be causal; “intermediate” means only variants with DAF in [0.0025, 0.0125) could be causal.

**Figure 3:**
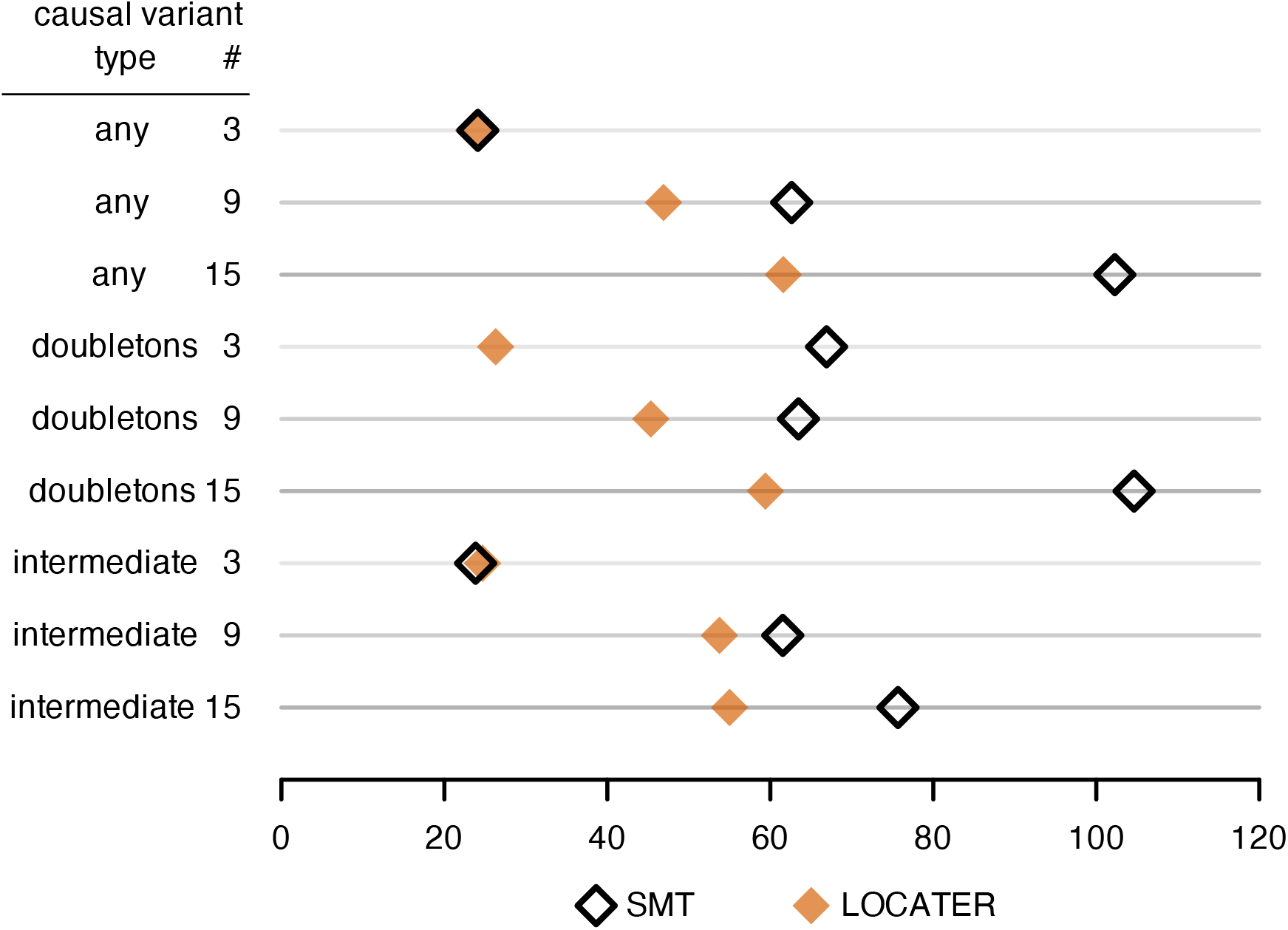
Dotplot of total association signal strength required to achieve 80% power (lower is better) under various simulation conditions where all causal variants were hidden.

From both Figure 2 and Figure 3 we see that LOCATER can reliably detect substantially weaker association signals than SMT across a variety of causal variant settings, especially when there are 9 or 15 causal variants. The only exception we see to this pattern is the case of 3 intermediate variants (derived allele count in [150, 750)), where SMT has a very slight advantage over LOCATER. The causal variants in the intermediate case are far too common to be picked up by SD, so LOCATER must rely on quadratic form testing to gain any advantage over SMT. This highlights the fact that quadratic forms struggle to have power against very sparse signals (e.g., only 3 causal variants).

Comparing Figure 3 to Figure 2, we see that the relative power gains available from LOCATER are typically less in the case of hidden causal variants compared to the case of observed causal variants, but still substantial, across settings. Except for the case of 3 doubletons, the power results reported in Figure 3 for SMT are remarkably similar to those reported in Figure 2 despite all of the causal variants being hidden. For the case of 3 doubletons, we see that the power of SMT is markedly reduced when the causal variants are hidden, making the relative power gain from LOCATER markedly large.

In order to confirm that these power results are robust to our choice of 10 kb as the size of the causal region, we replicated all of our experiments involving 9 causal variants assuming a 100 kb causal region. As can be seen in Section 5.4 (Supplementary Figures 28,29), the resulting power curves are very similar.

In order to compare LOCATER to the results one might obtain using sliding windows, we ran ACAT-O (STAAR without variant annotations) on our observed variant simulations from Figure 2, where any variant could be causal. Rather than testing all sliding windows for every simulation and effect size, we gave ACAT-O the precise location and width of the 10 kb causal window for each simulated dataset. This is an upper bound on the performance of ACAT-O in real-world settings where the location and size of the causal window are unknown. We ran ACAT-O in two different ways: one in which we restricted the variants considered to rare variants (MAF < 0.01) and another where all variants are tested regardless of frequency. As can be seen from Figure 4, the performance of both oracle ACAT-O approaches is roughly the same as SMT. LOCATER maintains its power advantage.

**Figure 4:**
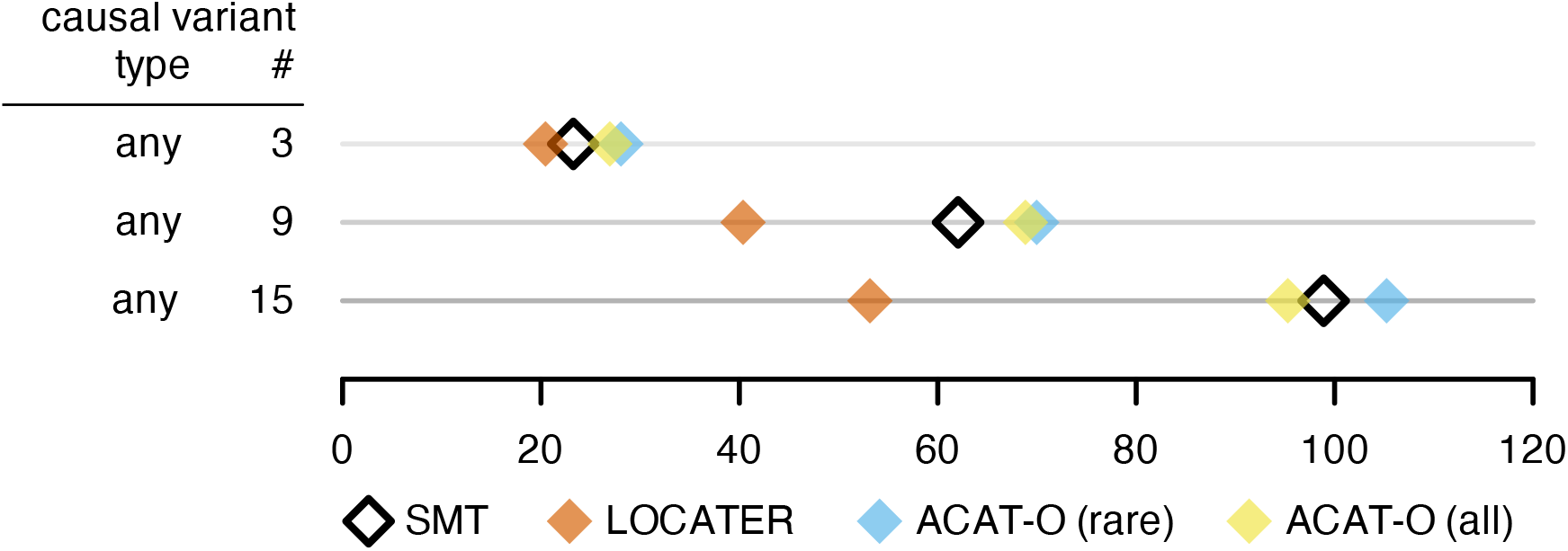
Dotplot of total association signal strength required to achieve 80% power (lower is better) under various simulation conditions where all causal variants were observed, including comparison to oracle ACAT-O methods that are given the causal variant window. ACAT-O (rare) only tests variants with MAF < 0.01 whereas ACAT-O (all) tests all variants within the causal window.

In Section 5.3, we pair each plot of power curves with a companion plot showing which LOCATER sub-test is driving the gain in power (Supplementary Figures 11,17,23,14,20,26). These results show that SD is typically the source of LOCATER power gains, not QForm, reflecting the statistical advantage of SD-based methods over quadratic form based procedures in the case of sparse signals [21]. While these simulations appear to imply that SD is only effective at capturing signals driven by very rare variants, this is expected since we only encoded very rare variants in the clade genotype matrix *X*^(*ℓ*)^ passed to SD. If SD was used to test the entire local genealogy at every locus, we may see increased statistical efficiency in incorporating common variant associations. We return to this point in the Discussion section. Substantial power gains are possible via LOCATER in the case of multiple rare causal variants.

### 2.6 Localization

Alongside power, an important factor for real-world utility is the ability of an association method to accurately localize causal variant(s) within a relatively narrow genomic interval. In Section 5.3, we also pair each plot of power curves with a companion localization plot (Supplementary Figures 12,18,24,15,21,27). To measure the ability of a given method to localize the causal region, we calculated the distance between the most significant marker (lead variant) reported by a method and the midpoint of the causal region in every simulation. We used these distances to estimate the width of an 80% confidence interval. This width represents the answer to the question: how large of a search window centered on the lead variant would an investigator need in order for that window to capture the midpoint of the causal region 80% of the time? In our plots reporting these confidence interval widths, we only report confidence interval widths at signal strengths where the corresponding method had at least 80% power to detect the causal region. As expected, both LOCATER and SMT struggled to localize the causal region more when the causal variants were hidden.

Across simulations where all variants in the causal region were potential causal variants, LOCATER more accurately localized the causal region than SMT, regardless of whether the causal variants were observed or hidden. This pattern also held when all of the causal variants were rare, the sole exception being the case of 3 hidden casual variants where SMT and LOCATER were effectively tied. Simulations where all of the causal variants were doubletons yielded somewhat more mixed results. With the exception of a few signal strengths when there were 9 causal variants, LOCATER still outperformed SMT in localizing the causal region when doubleton causal variants were observed. In all simulations where the doubleton causal variants were hidden, regardless of the number of causal variants, both SMT and LOCATER performed very poorly in localization, with both methods reporting confidence intervals wider than 600 kb across all signal strengths where they achieved at least 80% power. Overall our results suggest that LOCATER can leverage allelic heterogeneity to improve the localization of trait mapping compared to standard SMT.

### 2.7 Scalability

The ability to scale to large modern datasets with hundreds of thousands of samples is essential for the success of any trait mapping approach and the size of local genealogies presents a significant challenge. Based on our simulations involving 30k samples (60k haplotypes), LOCATER took an average of 19.14 seconds (sd: 2.88 seconds) to perform sprig testing and an average of 3.07 minutes (sd: 0.98 minutes) to perform quadratic form testing at each variant. These simulations were run on a shared-time university HPC cluster with heterogeneous nodes hosting a mix of CPU architectures. All jobs requested 8 cores and 160 GB of memory. When combined with the computational overhead required to form the clade genotype matrix *X*^(*ℓ*)^ and Ω^(*ℓ*)^ provided to LOCATER, our simulations required an average of 4.00 minutes (SD: 0.99 minutes) to test each target locus. When this is combined with the additional cost of performing ancestry inference with `kalis`, our simulations required an overall average of 6.42 minutes per target locus. These results suggest that future applications of LOCATER that only use inferred clades *X*^(*ℓ*)^ and avoid use of Ω^(*ℓ*)^ will achieve substantial computational savings.

In separate work, we also ran LOCATER in combination with `kalis v2` on a real sequencing dataset including 6795 individuals and 101 correlated quantitative traits (manuscript in prep.). We divided the genome into 4580 (partially overlapping) segments; each segment had an average of 13,000 variants. We allocated 12 cores and 60 GB of memory per segment. This allowed `kalis v2` to store two checkpoints in memory. The average time required for `kalis v2` and LOCATER to screen each segment was 32 minutes. That is equivalent to 3.35 years of single-core compute time. While substantial, this equates to 1.22 days using a cluster of 1000 CPUs. This result shows that it is feasible to run LOCATER on a moderately large genomic dataset using an academic compute cluster.

## 3 Discussion

We have presented a general framework for using inferred local ancestries to boost SMT association signals in the presence of allelic heterogeneity. To our knowledge, this is the first demonstration of any ancestry-testing approach that yields significant power gains over SMT in a genome-wide screen that includes non-coding regions. More importantly, our approach can be applied in conjunction with any ancestry inference engine, thus providing a flexible association testing framework that can adapt to rapidly improving ancestry inference methods.

In parallel work, we have demonstrated the efficacy of LOCATER in a dataset of 6,795 Finnish genomes with extensive quantitative trait data (manuscript in prep.), where we observed a significant power boost at several loci marked by allelic heterogeneity, and in our preliminary work we have run LOCATER on as many as 12,964 genomes on commodity hardware, which required 8,257 CPU-days to analyze 4 traits (unpublished data).

The largest power gains demonstrated in this paper were seen in the case of multiple rare causal variants evaluated using the SD subtest. This suggests that if we had tested more of the underlying tree structure at each locus with SD we may have achieved even greater power. In other words, as mentioned above, we may have clustered each distance matrix and tested more common clades than simply the sprigs with SD. This approach would present more of the underlying ancestral tree to LOCATER via *X*^(*ℓ*)^ rather than via Ω^(*ℓ*)^. Exploring the power of LOCATER at different points along the continuum between testing all of the ancestral structure with *X*^(*ℓ*)^ and testing all of the ancestral structure with Ω^(*ℓ*)^, will be a focus of future research.

In conjunction with the new features added to `kalis` [15], LOCATER provides an efficient method for genome-wide testing that is ready for use on real-world datasets now. These new features involve several algorithms — a general HMM checkpointing algorithm, a fast clustering algorithm, and a fast trace calculation method — that will likely prove helpful for the acceleration of other ancestry inference and association methods. Our novel quadratic form tail approximation approach, based on a shifted difference of chi-square random variables, provides a basis for emerging association methods to reliably test local relatedness matrices that may not be positive semi-definite.

Adequately adjusting for population structure when testing inferred local ancestries is an open and challenging problem. In this initial version of LOCATER, we allow principal components (PCs) to be included in *A*. As mentioned in Section 2.3, we also parameterized our novel quadratic form tail approximation in a way that naturally accommodates genomic-control-like inflation adjustments without requiring recalculating the genealogy at any LOCATER target loci. See Section 6.3 for details and theoretical motivation. However, future work applying LOCATER or any ancestry testing method to real genomic data will need to take special care when examining Q-Q plots for inflation.

LOCATER makes a number of critical methodological advances towards powerful ancestry-based association testing. We expect that further work building on these advances alongside the application of LOCATER to more diverse datasets will yield new functional discoveries.

## 4 Methods

### 4.1 Haplotype Data Simulation

In order to assess the calibration and power of LOCATER, we simulated 100 genomic datasets, each consisting of a 1 Mb chromosome for 30k human samples (60k haplotypes). Each dataset was simulated using `msprime` [36]. In order to model the diversity of arising genomic datasets, 10k samples in each dataset were drawn from each of three 1000 Genomes populations – Yoruba, Han Chinese, and Central European. See Section 6.5 for further details.

### 4.2 Phenotype Simulation

In the middle of each 1 Mb region, we selected causal variants within a 10 kb causal window. We fully replicated these simulations under all 18 possible combinations of 3 parameters: the number of causal variants, the allele frequency constraint imposed on those causal variants, and whether the causal variants were assumed to have been observed (called during sequencing) or hidden. More explicitly, we considered the case of 3, 9, or 15 causal variants. These causal variants were selected uniformly at random from among variants within the 10 kb causal window meeting the given allele frequency constraint. As our primary focus, we considered the case of no allele frequency restraint, in which case every variant in the 10 kb causal window had an equal chance of being selected as a causal variant. We also considered the case where all causal variants were constrained to be doubletons (present in two copies) and the case where all of the causal variants were constrained to have derived allele frequency in the half open interval [0.0025, 0.0125). If the simulated chromosomes did not include the requisite number of causal variants within the 10 kb causal window, we rejected that simulated dataset and simulated a new set of chromosomes.

Given an active set of causal variants, we simulated *Y* while distributing the observed effects across the causal variants as evenly as possible by manipulating the *QR*-decomposition as done by [21]. Following their approach, let 𝒜 denote this selected set of causal variants and *X*_*𝒜*_ denote the genotype matrix encoding those causal variants. Consider the *QR*-decomposition 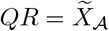, which we define as 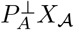 with length-normalized columns. The sufficient statistic for the oracle model is 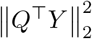 with expected value 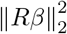. For a desired total association signal strength *s*, we solve 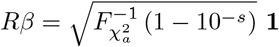 to make the magnitude of each entry of *β* as similar as possible. Then, we simulate 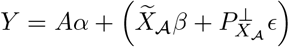 where *ϵ* ∼ *N* (0, *I*_*n*_). This ensures that the observed 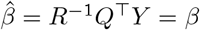 is stable across simulations and that the − log_10_ p-value that one would obtain by testing the resulting *Y* with an oracle ANOVA model that “knows” the active predictors will be approximately *s* in every simulation.

### 4.3 Checkpointing

Let *j* index the positions of the *L* target loci from smallest to largest. We will solve for an optimal checkpointing schedule for the forward algorithm to propagate to target loci sequentially in reverse order, from *j* = *L* to *j* = 1. We will use *j* = 0 to index the HMM prior hidden state probabilities used to initialize the forward algorithm. Informally, for each subsequent target locus, a checkpointing schedule states the checkpoint at which the forward algorithm should start and the intervening target loci where it should stop to store any new checkpoints on its way to reach that target locus. To optimize this schedule, we do not need to consider storing checkpoints at variants that are not target loci since any such checkpointing strategy could be improved by moving those checkpoints to target loci. Thus, our checkpointing schedule will only need to render integers in 1, …, *L*.

Optimizing the checkpointing schedule is based on recycling the memory used to store checkpoints that are no longer being used. Here, we solve for an optimal checkpointing schedule where the computational cost to propagate the forward algorithm from any position to any other may be arbitrary. The checkpointing routine that minimizes the total computational cost in this setting may be obtained by solving nested optimization problems. While tractable, solving this system of nested optimization problems can be rather time consuming to solve. Therefore, we subsequently consider the special case where the cost of propagating the forward algorithm between consecutive target positions is constant. For example, in a genomics context this may be true if we have equally spaced target loci. In that special case, we explain how the optimal checkpointing schedule can be solved rapidly via dynamic programming.

We start with the general case. Define *g* : 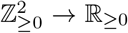 such that *g* (*s, t*) is the cost of propagating the forward algorithm directly from target locus *s* to target locus *t*, defined to be zero whenever *s* = *t*. Define 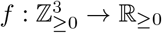 such that *f* (*s, t, c*) is the minimal cost required for the forward algorithm to sequentially visit target loci *t, t* − 1, *t* − 2, …, *s* given *c* available checkpoints and a known set of forward probabilities (i.e. a checkpoint, or the HMM prior) at *s*. That is, *f* (*s, t, c*) corresponds to the cost of the (unknown) optimal schedule given memory for *c* checkpoints.

We aim to obtain a checkpoint schedule which enables us to achieve the cost *f* (0, *T, C*). This objective is defined by the three following equations. First, if there are no checkpoints available, simply propagating to each target locus sequentially from the initial position is our only option, which comes with cost 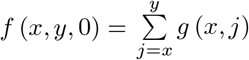. Second, if we already know the forward probabilities at a given position, there is no computational cost to obtaining them, so *f* (*x, x, c*) = 0 for any *c*. Third, we have the following recursive relationship

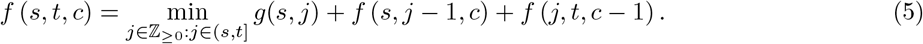

Intuitively, this recursion expresses the idea that solving for an optimal checkpointing schedule with *c* checkpoints over an interval can be thought of as placing one of the checkpoints at some optimal index to divide the interval so that the upper part is solved with *c* − 1 checkpoints; the lower part, *c* checkpoints. We will write that optimal index, the argument that achieves the minimum of Equation (5) as

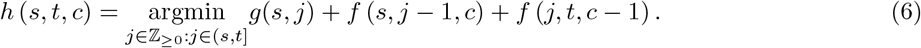

Together, these three equations allow us to solve for an optimal checkpoint schedule by transversing a bifurcating tree from right to left, from the leaves toward the root, where each node can be identified with an optimal index *h*.

While solving this recursion is tractable, it can be rapidly accelerated if we assume that the computational cost of propagating the forward probabilities between adjacent target loci is fixed. In other words, we assume *g* (*x* − 1, *x*) is constant for all target indices *x* ∈ {1, 2, …, *L*}. This makes the computational cost a simple linear function of the distance between indices, yielding the following simplified system of equations.

Analogous to above, we have 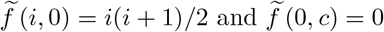 for all *c*. Our recursions simplify to

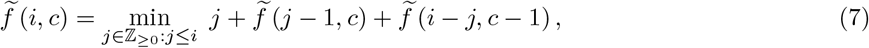

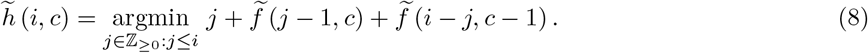

For some pre-specified maximum number of target loci *L* and maximum number of available checkpoints *C*, let us define two matrices that we will use as look-up tables for our dynamic program. Let 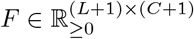 such that 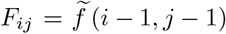 and 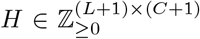 such that 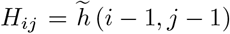. Given the nested nature of 7 and 8, we can rapidly solve for each entry of *F* and *H* by synchronously iterating over the entries of both matrices in column-major order. Once we have a completed *F* and *H*, we can use 8 to read off the appropriate entries in table *H* to obtain the optimal checkpointing schedule for any problem with up to *L* target variants and *C* available checkpoints. Methods for constructing *H* and reading it to construct a checkpoint schedule are available in `kalis v2`. This implementation covers the slightly more general case than (7) where the cost depends on distance but is independent of locus. This corresponds to a translation invariance assumption that does not cover the most general case in (5).

### 4.4 Defining our Generalized Relatedness Matrix

Here we build up to our definition of a generalized eGRM, which we will pass to LOCATER as Ω^(*ℓ*)^. We will construct Ω^(*ℓ*)^ based on an asymmetric genetic distance matrix 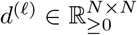, such as the one provided by `kalis` (see Equation (2)), and a set of monotonic regularization functions *g*_1_, …, *g*_*N*_ which we will introduce shortly. Recall that 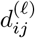 measures the distance to haplotype *i* from haplotype *j*. The distance from any given haplotype to itself 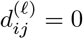. Let *π*_*j*_ : [*N*] → [*N*] be the permutation that sorts 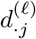 such that 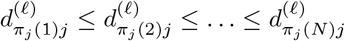. By convention, *π*_*j*_(1) = *j* and *π*_*j*_(*N* + 1) = *π*_*j*_(*N*). Using *d*^(*ℓ*)^, we define a local haplotype relatedness matrix 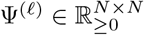 with elements

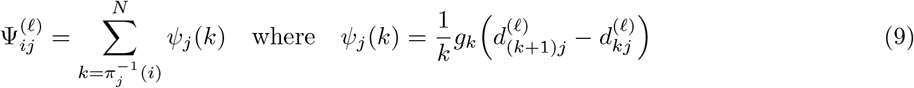

and each *g*_*k*_ : ℝ_≥0_ → ℝ_≥0_ is a monotonic function of *x* such that *g*_*k*_(0) = 0. Note, given these definitions, *ψ*_*j*_(*N*) = 0.

Our construction does not require the assumption of diploid samples but we will assume that here for ease of exposition. We will assume that the rows and columns of Ψ^(*ℓ*)^ are permuted such that haplotypes from the same sample are grouped together. This allows us to succinctly write our generalized eGRM in terms of Equation (9) as

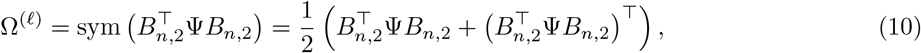

where *B*_*n*,2_ = *I*_*n*×*n*_ ⊗ **1**_2_ and ⊗ is the Kronecker product.

In Section 6.2 we explicitly show how this construction of Ω^(*ℓ*)^ generalizes the standard eGRM. In short, there we show that the eGRM can be expressed in terms of a haplotype similarity matrix assuming Hardy–Weinberg equilibrium and specific choices for the background covariates and allele frequency weights. Then we connect that haplotype similarity matrix representation to our choice of Ψ^(*ℓ*)^ in Equation (9) above.

### 4.5 Efficiently Constructing Clade Genotypes and our Generalized eGRM in LOCATER

Building on the notation above in Section 4.4, currently in LOCATER and this paper, we set

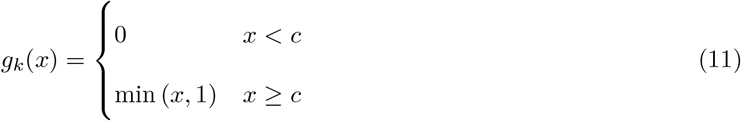

for all *k* where our threshold *c* = −0.2 log (*μ*) and *μ* is the mutation probability parameter provided to the LS model. This choice of regularization function(s) tends to filter out many low evidence clades. This function also restricts the clade matrix representation so that at most one mutation can be present on a given branch.

Given such a regularization, Ψ_·*j*_ has a series of nested neighborhoods of donor haplotypes along *i* = *π*_*j*_(1), *π*_*j*_(2), …, *π*_*j*_(*N*) where there are distances of at least *c* between adjacent level sets of distances. This allows us to represent the level sets of Ψ_*ij*_ as the solution to a clustering problem on the real interval from 0 to the maximum possible distance (*D*) where we require unique clusters to be at least distance *c* apart. Here, each cluster corresponds to a level set of donor haplotypes. We use a single-pass partial sorting algorithm based on doubly-linked-lists to solve this clustering problem in 𝒪(*N*) time. In our experiments on simulated haplotype data, our partial sorting algorithm achieves roughly an order of magnitude speed up over merge sort. Given the definition of *υ* in Equation (2), the maximum possible distance is *D* = − log(*υ*) ≈ 744.44. This maximum is helpful in accelerating our implementation because the number of possible level sets (clusters) *d* is bounded above *d* ≤ ⌈*D/c* + 1⌉, allowing us to efficiently pre-allocate sufficient memory to store the clustering solution. Since this partial sorting algorithm can be run in parallel for each recipient haplotype (for each column Ψ_·*j*_), we use a multi-threaded implementation in `kalis v2`.

Let 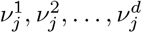 denote the nested neighborhoods corresponding to recipient haplotype *j* ordered from the nearest neighborhood to the furthest possible neighborhood. Some of the more distant neighborhoods may be empty if fewer than *d* clusters (level sets) are observed among the distances 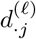. From these neighborhoods, we call a set of clades for each layer 1 through *d* by iteratively identifying fully connected cliques within the directed graph implied by the neighborhoods as follows. In this single-pass clustering algorithm, we also compute the level sets of Ψ_*ij*_, the nearest-neighbors of *j* and various other quantities so that the downstream modifications we make to Ψ related to sprig testing may be done in place (without recalculating Ψ). Many pairs of columns of Ψ are then calculated in parallel. By processing columns of Ψ in pairs, we can directly compute our sample by sample matrix 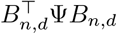 in a single pass. Symmetrizing this matrix yields Ω^(*ℓ*)^ as defined in Equation (10). Future work will focus on reducing the computational and memory requirements of this symmetrization step.

### 4.6 Parallelized Stable Distillation Procedure

Given a matrix of inferred, clade-based genotypes *X*^(*ℓ*)^, we use the one-predictor-at-a-time SD procedure described in Equation 4 of [21] equipped with the simple quantile filter presented in Algorithm 1 of [21]. In this SD procedure, we take (*A, G*^(*ℓ*)^) as the background covariates when testing 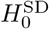 at a particular target locus *ℓ*. In short, using this approach, we “distill” one *β*_*j*_ for *j* = 1, …, *p* at a time, obtaining an independent p-value for each. These p-values are then tested using the Rényi Outlier Test [32]. To run this procedure, LOCATER requires an estimated upper bound *c* on the number of independent causal clades: *c* ≥ |{*j* : *β*_*j*_ ≠ 0}|. By default and throughout this paper, we set *c* = 16. This bound *c* is used to set the simple quantile filtering threshold used during distillation. Explicitly, LOCATER sets the quantile filtering threshold 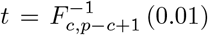 where *F*_*a,b*_ is the CDF of the Beta distribution with expectation 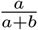. By default, the maximum number of outliers considered by the Rényi Outlier Test is set to *c*.

### 4.7 Quadratic Form Testing & Tail Approximation

Let *QR* = (*A, G*_*j*_) be the QR-decomposition of the *n* × (*q* + 1) matrix (*A, G*_*j*_) and let *P* = *I* − *QQ*^⊤^ project onto the subspace orthogonal to the columns of (*A, G*_*j*_). Differentiating the Gaussian likelihood corresponding to Equation (1) with respect to *τ* yields *Y*^⊤^*P*Ω^(*ℓ*)^*PY* as the score statistic. Under 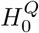,

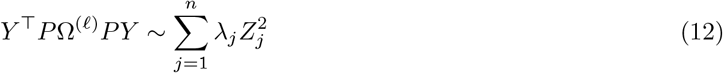

where each 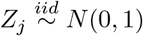. Following the approach of FastSKAT [34], we use partial eigendecomposition to obtain a computationally tractable approximation to this null distribution. Given, a top-*k* eigendecomposition in which we explicitly calculate the leading *k* eigenvalues, we have

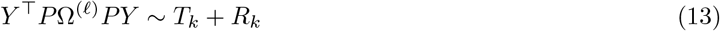

where 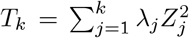and 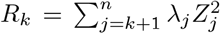. Since *λ*_*k*+1_, …, *λ*_*n*_ are unknown, FastSKAT proposes approximating the distribution of *R*_*k*_ using a single chi-square random variable 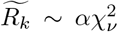. The scale parameter *α* and degrees of freedom parameter *ν* are set to match the mean and variance of *R*_*k*_, as initially proposed by Satterthwaite [33]:

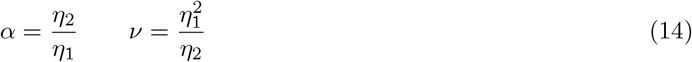

where 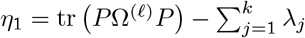 and 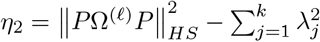. Substituting 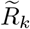 in for *R*_*k*_, FastSKAT uses 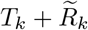 as an approximate null distribution. This is still a linear combination of chi-square random variables, but since all of its parameters are now known, its distribution is available via the fast Fourier transform.

Unfortunately, in the context of LOCATER, 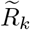 does not provide an adequate approximation to the distribution of *R*_*k*_. We found that the tails of 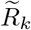 tended to decay faster than those of *R*_*k*_, yielding anti-conservative estimates of more extreme p-values. This is because the scale *α* in Equation (14) can over-estimate the rate of decay in the tail of *R*_*k*_. On a technical note, the Satterthwaite approximation also assumes that *P*Ω^(*ℓ*)^*P* is positive semi-definite (PSD). This is not at all guaranteed in our application. The closeness of any *P*Ω^(*ℓ*)^*P* to PSD will depend on the ancestry at *ℓ* as well as the user’s choice of ancestry inference engine and clade-encoding method. The *P*Ω^(*ℓ*)^*P* matrices we have observed in the development of LOCATER are typically close to, but not exactly, PSD.

In order to make the tails of our approximating distribution more robust (heavier) and obviate the PSD requirement, we propose approximating *R*_*k*_ with a difference of independent chi-square random variables:

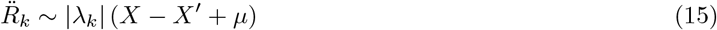

where 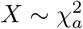 is independent of 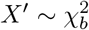. A derivation and explicit equations for the three scalar parameters – *a, b*, and *μ* – are given in Section 6.3 in terms of *η*_1_ and *η*_2_. In short, these parameters are set so that our approximation matches the mean and variance of *R*_*k*_ while simultaneously minimizing |*μ*|. As shown in Section 6.3, a key advantage of our specification of 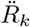 is that it can be easily generalized to accommodate three inflation parameters — *ν*, 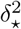, and 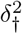 — that are orthogonal to *a, b*, and *μ*. There we show how these parameters naturally arise when considering the presence of some unobserved confounder. The orthogonality allows us to tune *ν*, 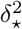, and 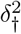 to adjust for population confounding after observing an initial Q-Q plot of p-values 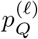 without having to recalculate *a, b*, or *μ*. In other words, we can adjust 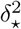, and 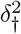 without having to re-calculate the genealogy at any target variant. While we do not make use of that capability in this paper, we expect it will play an important role in future analyses.

Critical to the scalability of this approach is the availability of *η*_1_ and *η*_2_ via fast trace calculations. FastSKAT [34] uses stochastic estimator of *η*_2_. Starting with a symmetric Ω^(*ℓ*)^, LOCATER employs a fully parallelizable and distributable trace calculation routine that obtains both *η*_1_ and *η*_2_ with fewer than 7(*q* + 1)*n*^2^ + 3*n*(*q* + 1)^2^ FLOPs. Recall that *q* is the number of columns in our background covariate matrix *A*. See Section 6.4 for further details.

### 4.8 Combining p-values: MSSE

The classic Fisher method [37] of combining a vector of p-values *U* = (*U*_1_, …, *U*_*p*_)^⊤^ is based on comparing the test statistic

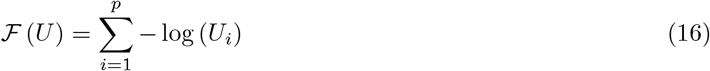

to its null distribution: a gamma distribution with scale parameter 1 and shape parameter *p*, whose survival function we denote by *G*_*p*_. Since it takes the form of simple sum, ℱ implicitly considers all possible alternatives: any combination of the p-values may be non-null. However, in applications such as LOCATER, where we are only interested in certain subsets of alternatives, this can lead to an unnecessary loss in power.

The aim of LOCATER is to enhance SMT signals by leveraging allelic heterogeneity. In a genomic region with allelic heterogeneity, we expect the most significant LOCATER signal to arise at a target locus where the core marker at that locus tags one of the causal alleles. Thus, we are only interested in testing combinations of p-values where *p*_SMT_ is non-null. This in effect reduces the number of combinations of non-null p-value alternatives we want to test, which we can achieve by using

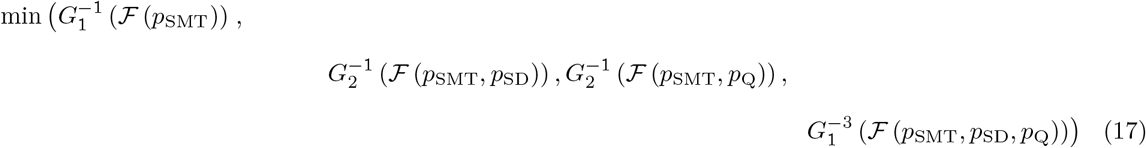

as our test statistic. The null distribution of this test statistic can be easily pre-calculated via simulation. Using these simulations, we summarize the body of the null distribution via a monotone cubic spline and fit the exponential tail of the null distribution to obtain a rapidly computable test.

## Acknowledgments

RC and XW were supported by NIH grants R01HG013371-01 and UM1HG008853 to IH. LJMA was partially supported by the EPSRC research grant “PINCODE”, reference EP/X028100/1, and UKRI grant, “OCEAN”, reference EP/Y014650/1. The authors would like to thank Nathan Stitziel, Chris Holmes, and Chris Spencer for helpful advice and discussions.

## 5 Supplementary Figures

### 5.1 Checkpointing

**Figure 5:**
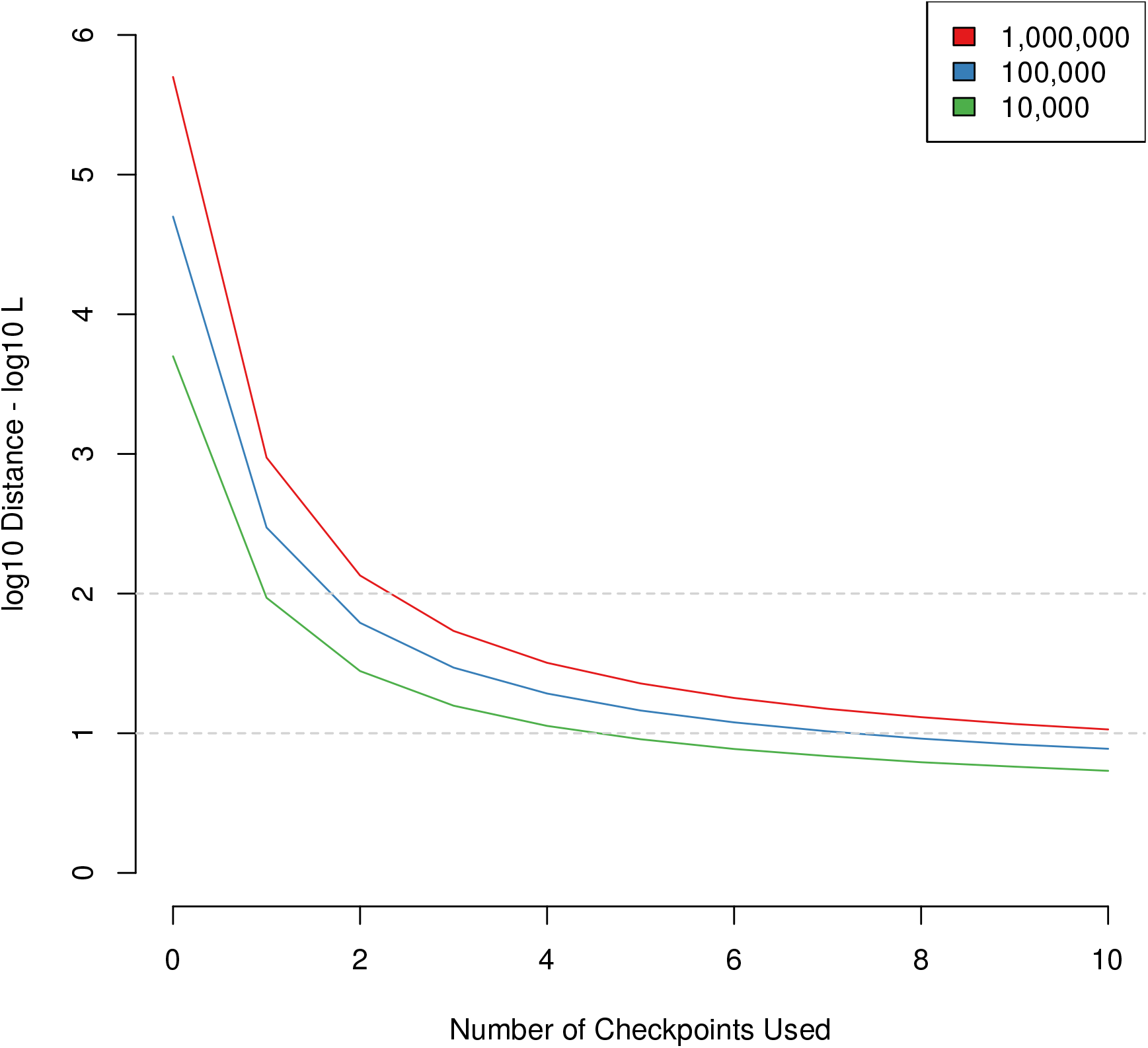
Computational Cost Using Optimal Checkpointing. Here we consider the total number of variants the forward algorithm needs to iterate over (the total distance *D* it has to cover) in order to visit all loci on a chromosome of length *L* using our optimal checkpointing strategy. Here we plot the log_10_ (*D/L*) as a function of the number of checkpoints, *C*, available for chromosomes of three potential lengths indicated by the legend: *L* ∈ {10^4^, 10^5^, 10^6^}. The dotted gray horizontal lines highlight that we are able to reduce *D* to under 100*L* with only 3 checkpoints and nearly 10*L* with only 10 checkpoints in all cases.

### 5.2 Null Simulation Q-Q Plots

**Figure 6:**
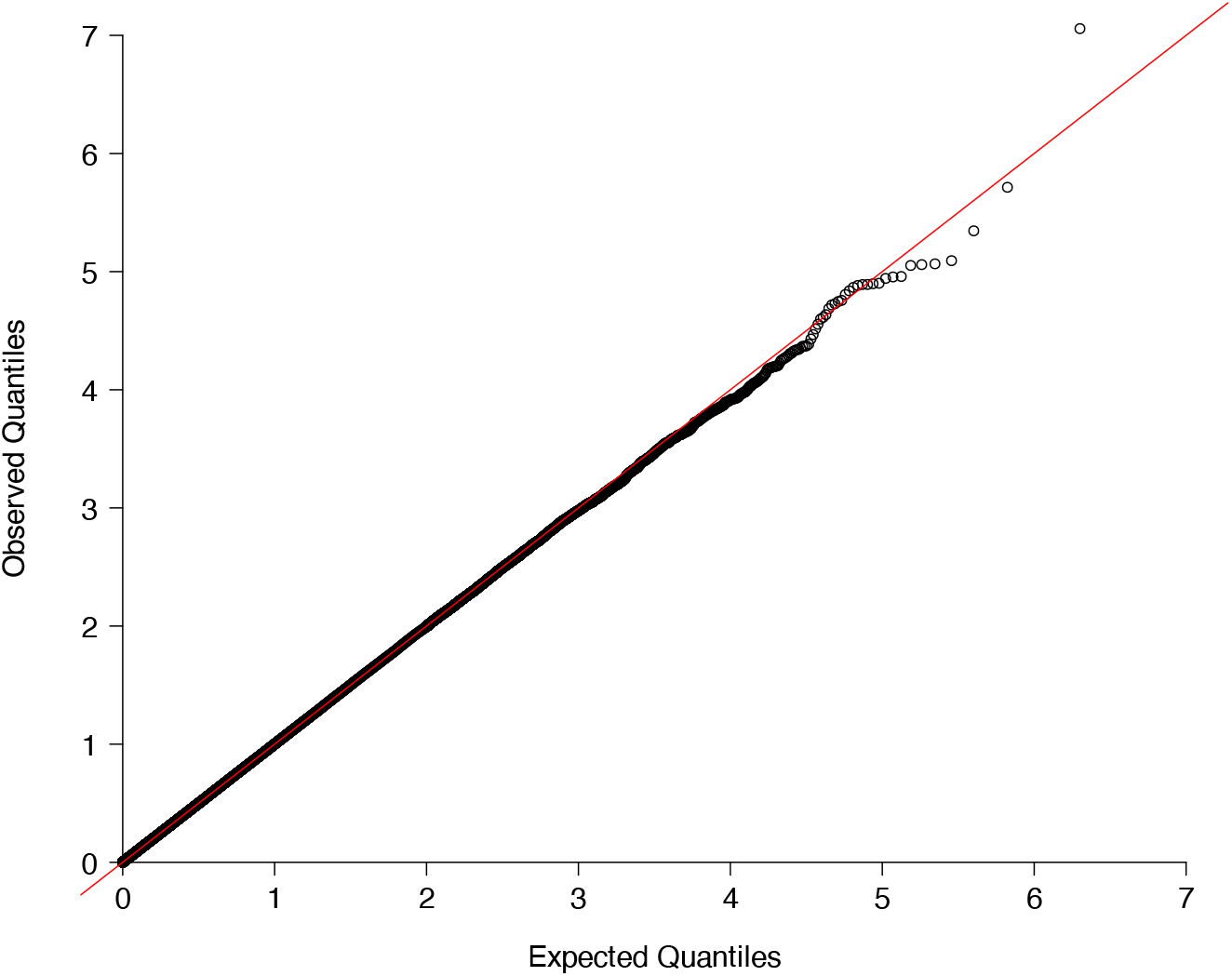
SMT Subtest Q-Q plot. Q-Q plot of samples of – 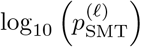 based on 1 million independent phenotype vectors simulated under the null hypothesis.

**Figure 7:**
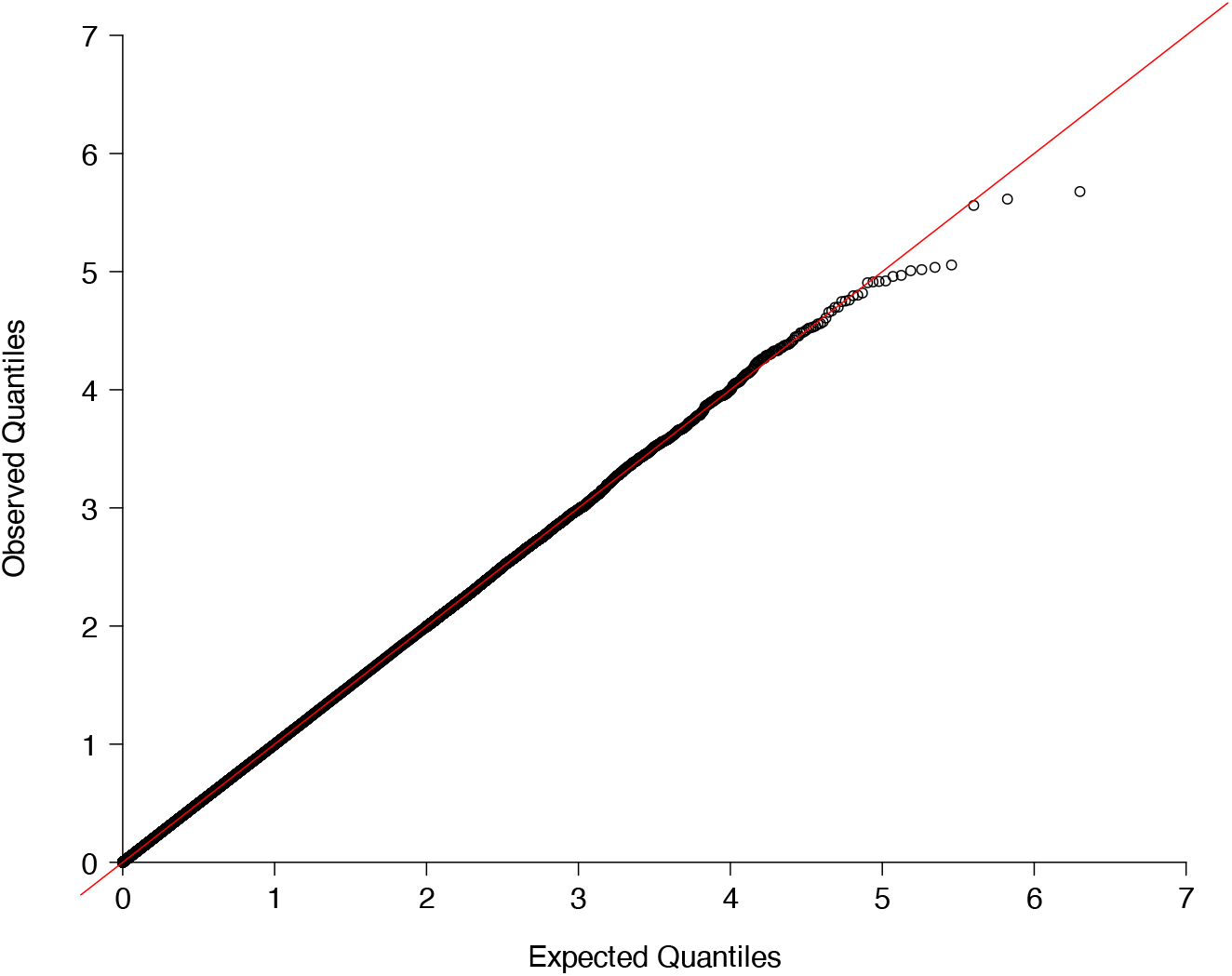
SD Subtest Q-Q plot. Q-Q plot of samples of – 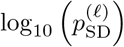 based on 1 million independent phenotype vectors simulated under the null hypothesis.

**Figure 8:**
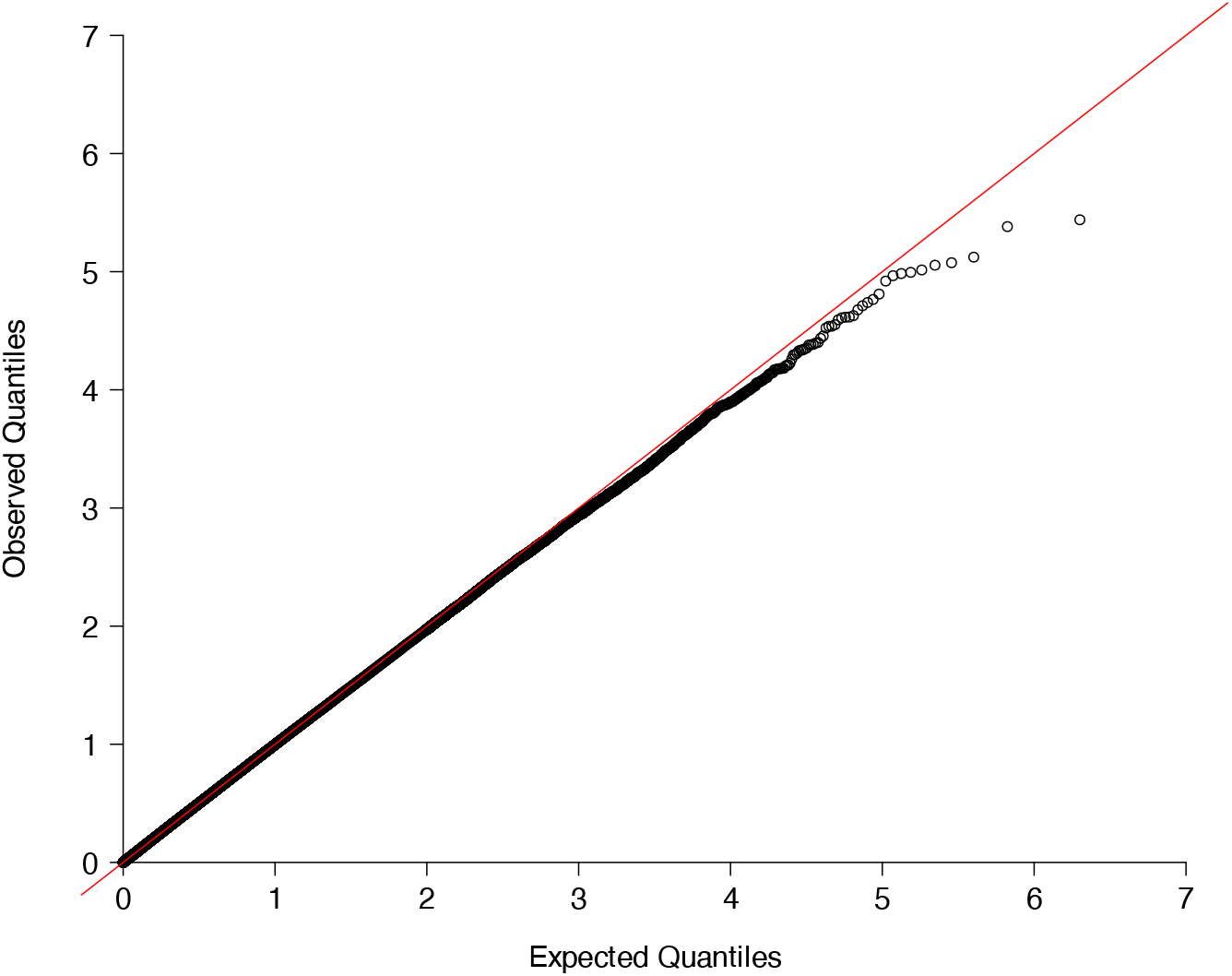
QForm Subtest Q-Q plot. Q-Q plot of samples of – 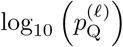 based on 1 million independent phenotype vectors simulated under the null hypothesis.

**Figure 9:**
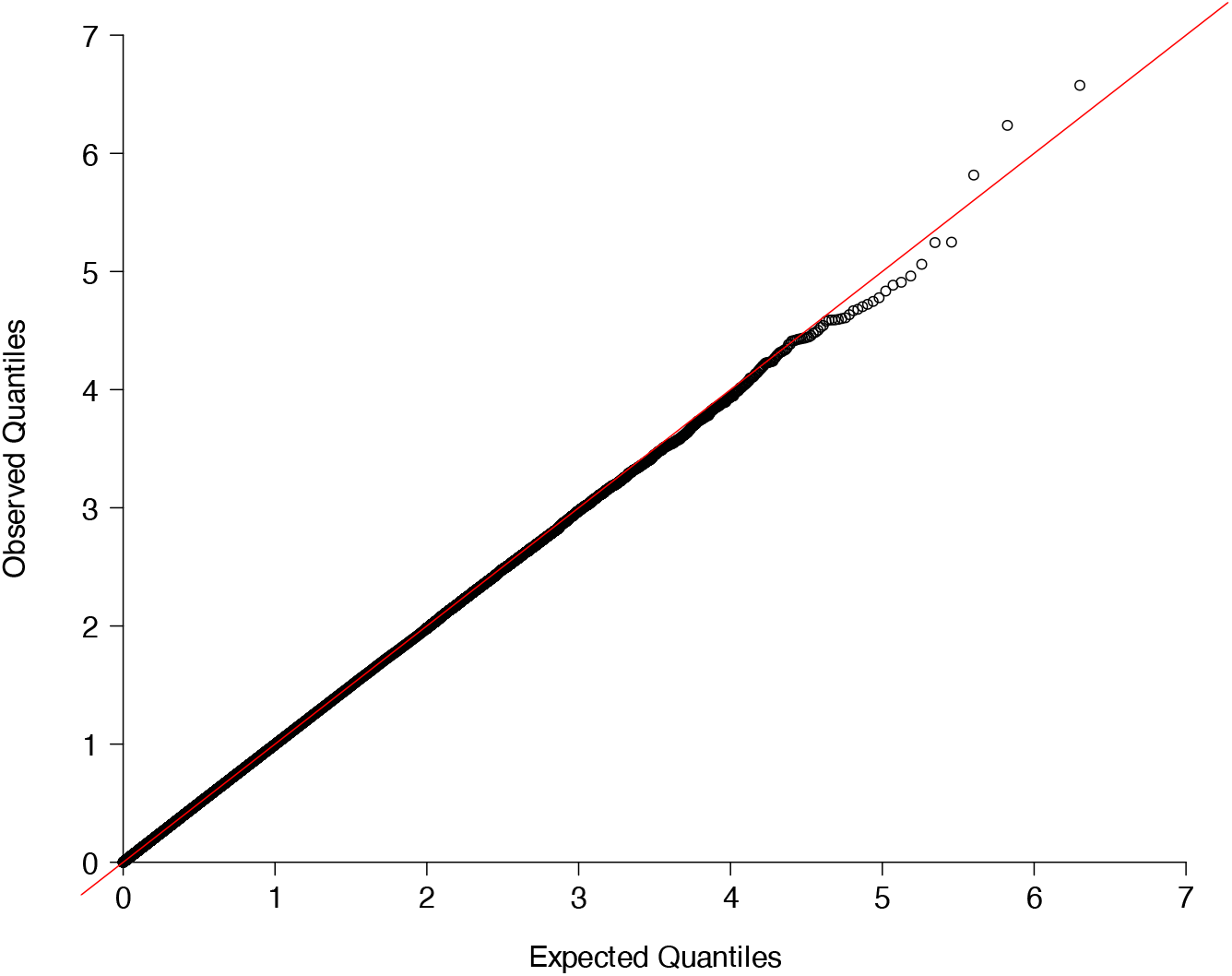
LOCATER Combined Q-Q plot. Q-Q plot of samples of – 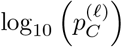 based on 1 million independent phenotype vectors simulated under the null hypothesis.

### 5.3 Power, Source, & Localization Curves

**Figure 10:**
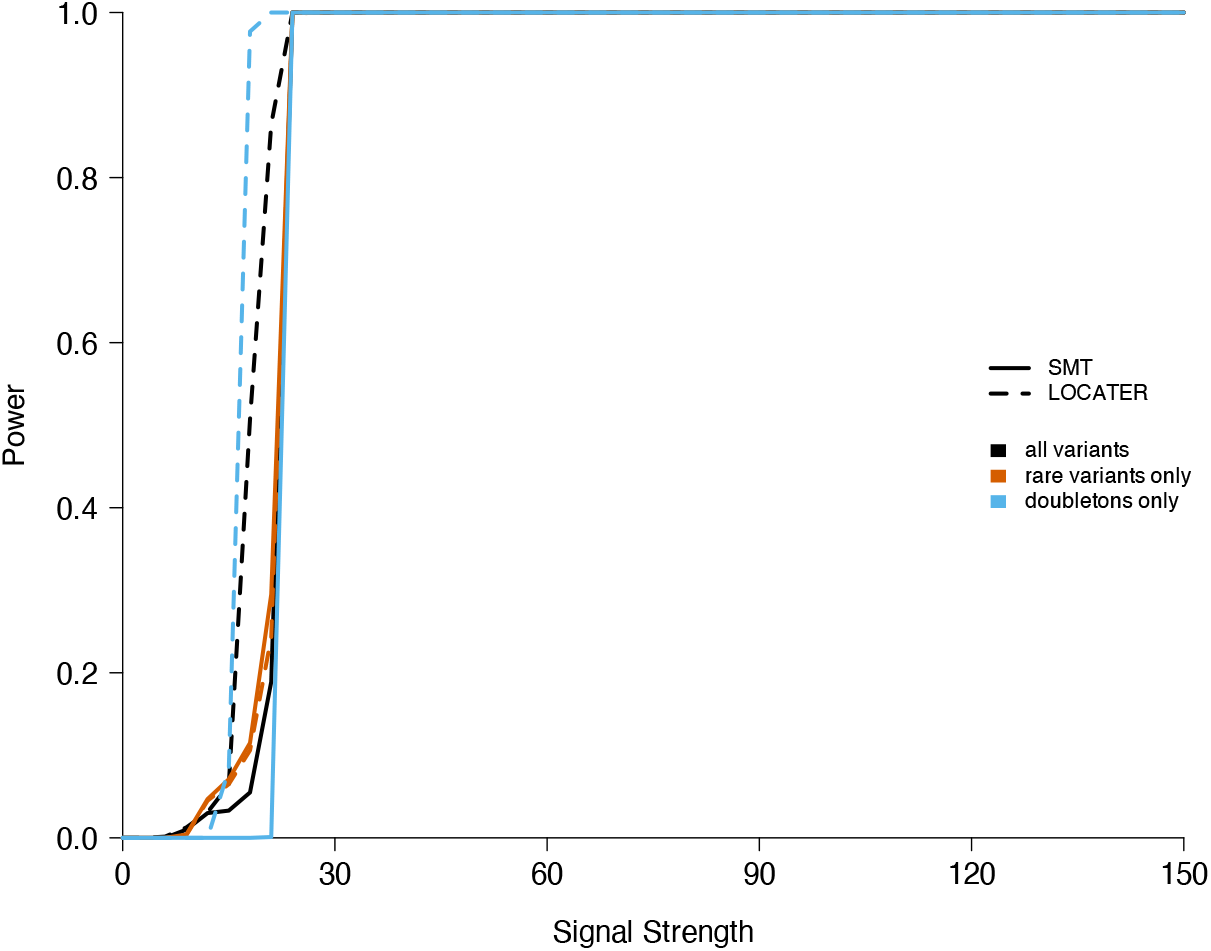
Power comparison between SMT and LOCATER in simulations where there are 3 causal variants, all observed.

**Figure 11:**
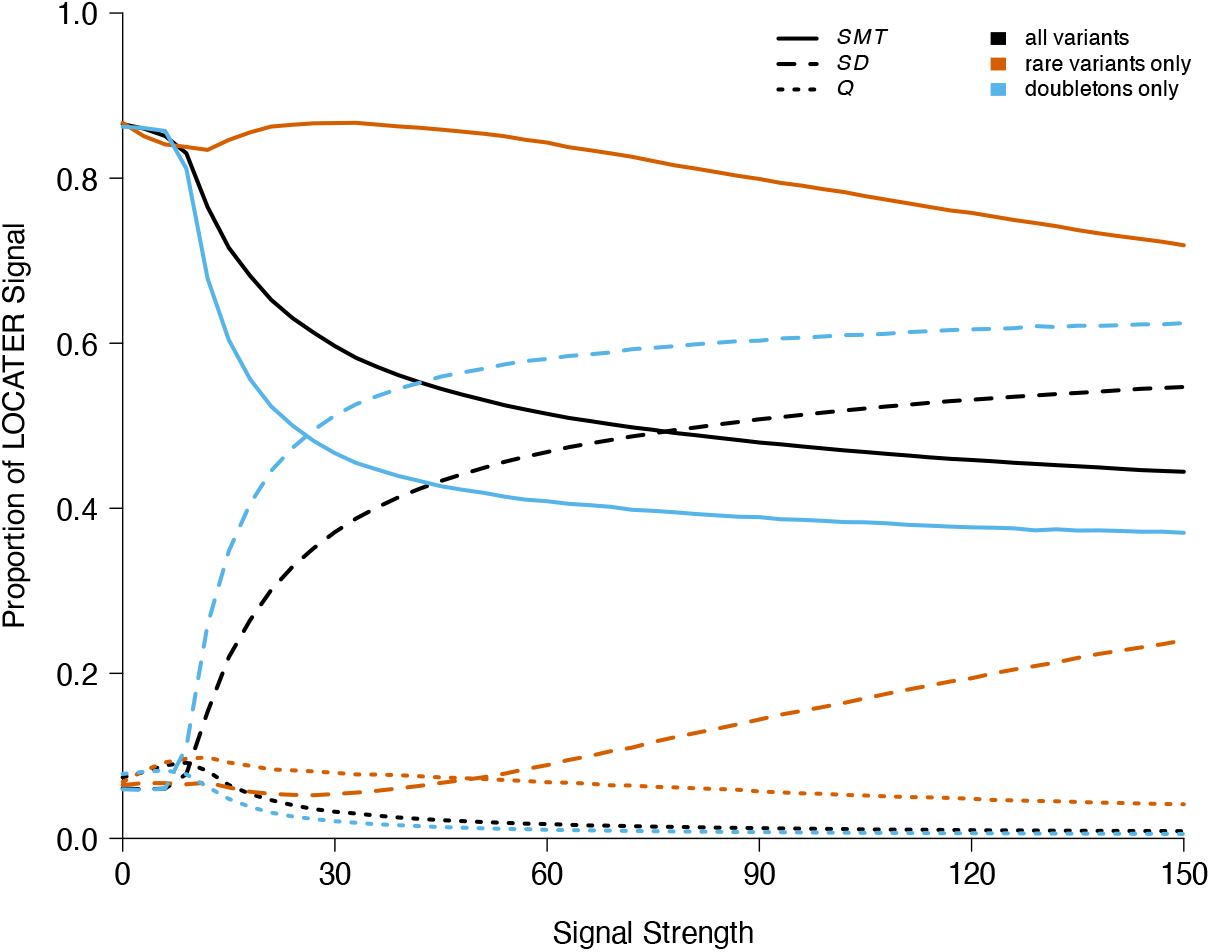
Average proportion of signal driven by each sub-test within LOCATER in simulations where there are 3 causal variants, all observed. For every simulation, the proportion of signal attributed to the *SMT* part of LOCATER (solid lines) defined as 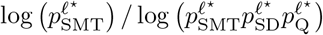 where *ℓ*^⋆^ is the locus *ℓ* that achieves the largest combined LOCATER p-value. The proportion of signal attributed to *SD* (dashed lines) or quadratic form testing, abbreviated as *Q* (dotted lines), is defined analogously.

**Figure 12:**
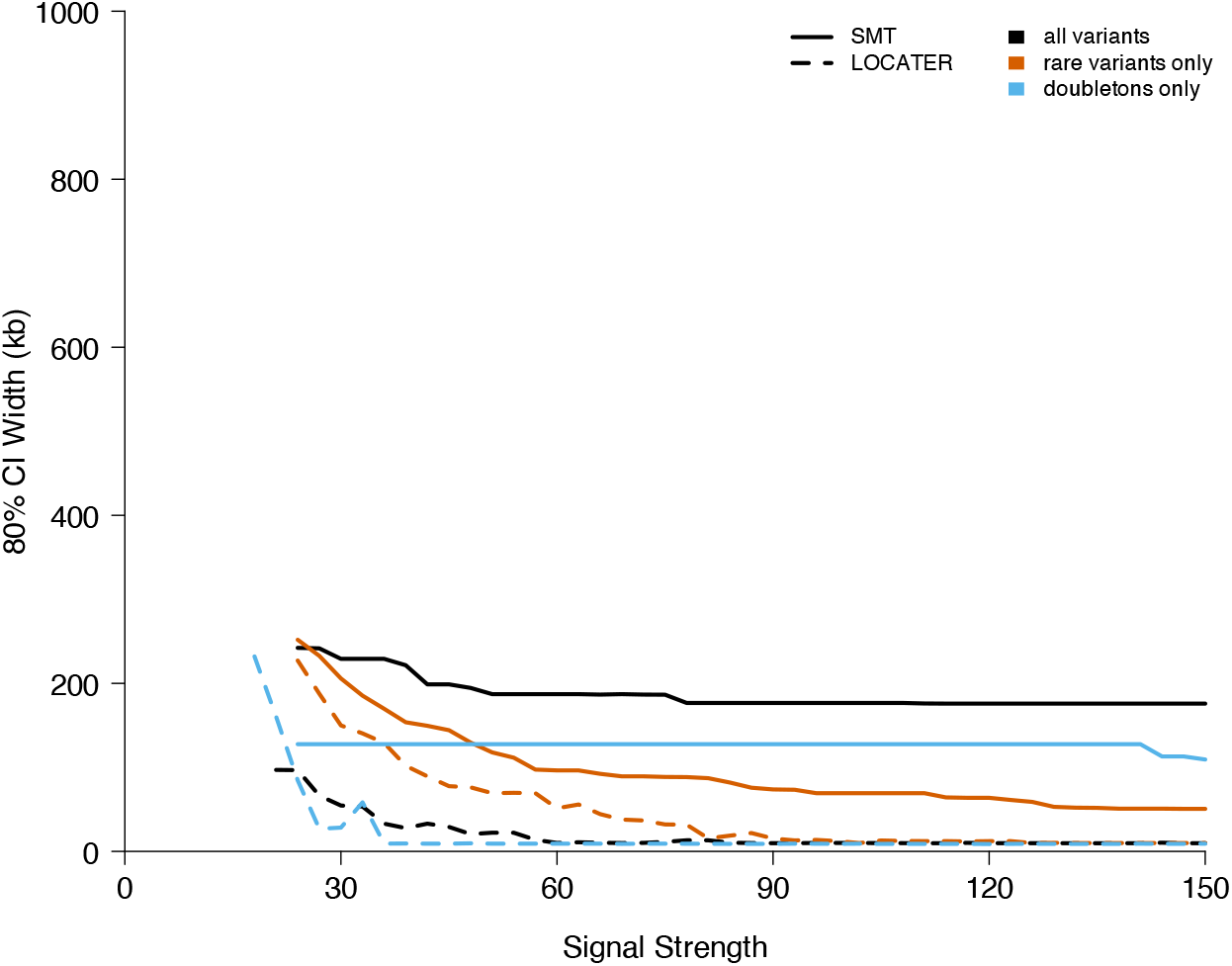
Width of confidence intervals (CI), centered on the position with the largest signal, that cover the midpoint of the causal region in 80% of simulations where there are 3 causal variants, all observed. In simulations where multiple variants are tied to have the largest association signal, we take the distance to the midpoint of the causal region to be the average distance from each of the tied variants.

**Figure 13:**
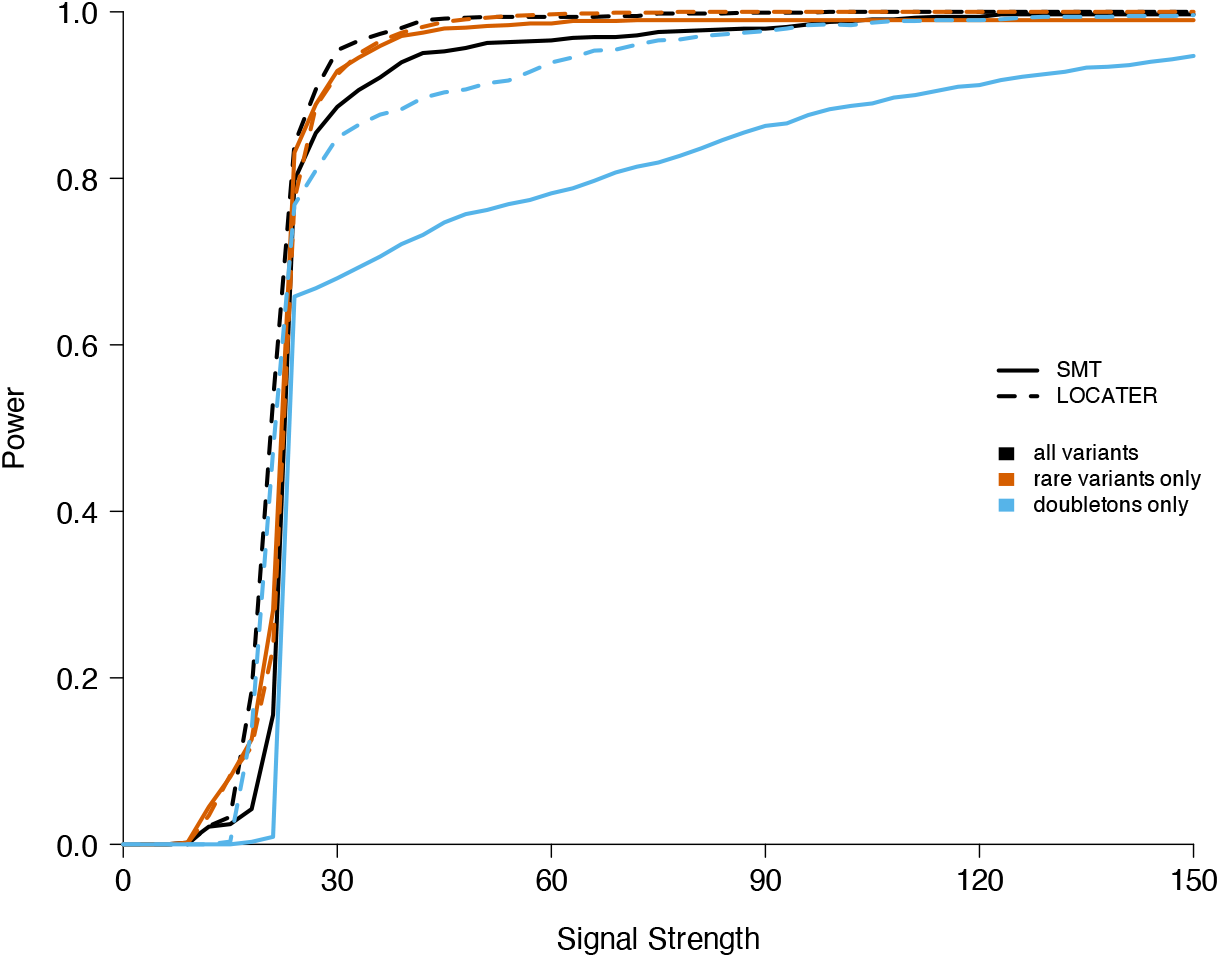
Power comparison between SMT and LOCATER in simulations where there are 3 causal variants, all hidden.

**Figure 14:**
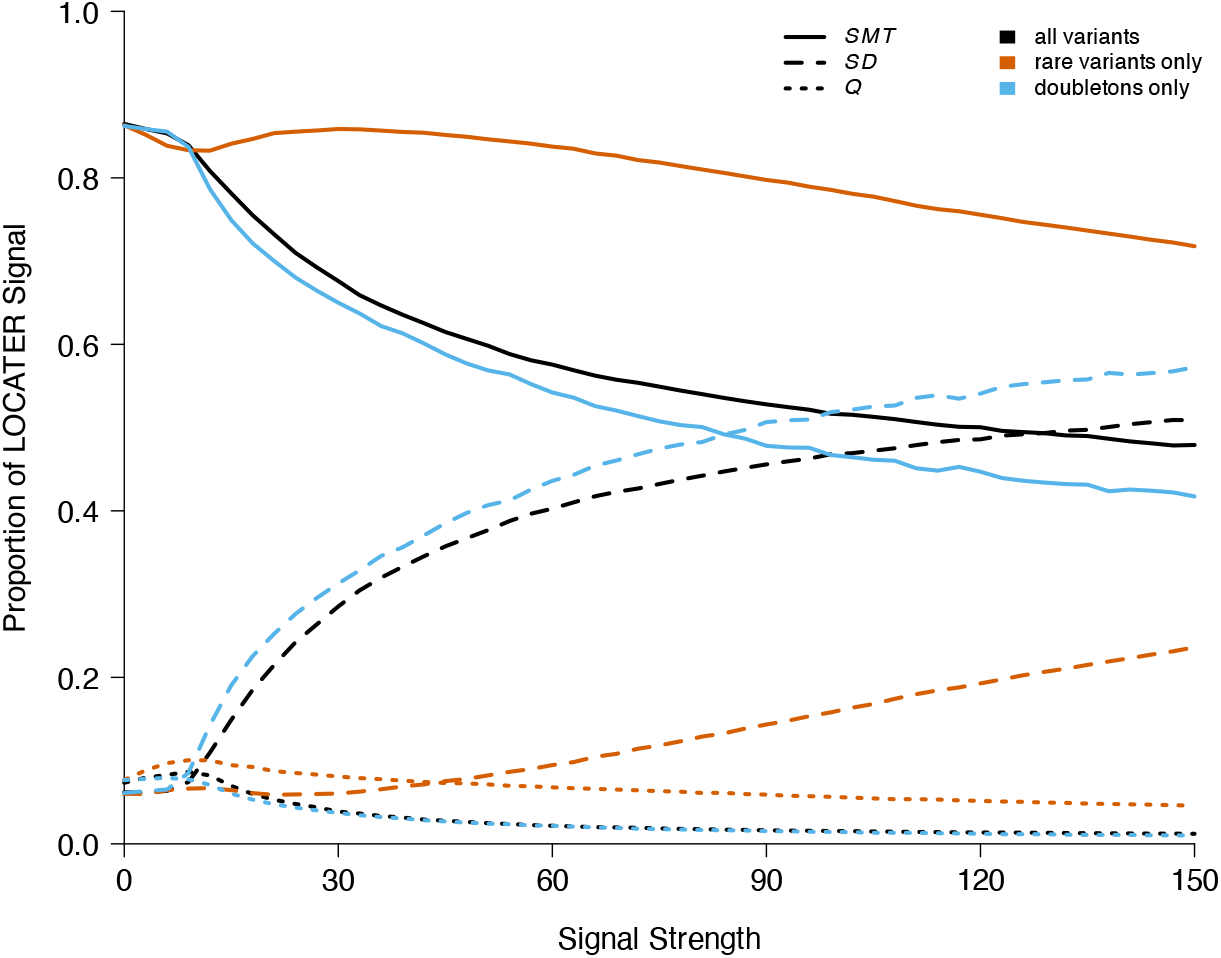
Average proportion of signal driven by each sub-test within LOCATER in simulations where there are 3 causal variants, all hidden. For every simulation, the proportion of signal attributed to the *SMT* part of LOCATER (solid lines) defined as 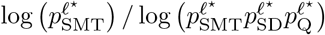 where *ℓ*^⋆^ is the locus *ℓ* that achieves the largest combined LOCATER p-value. The proportion of signal attributed to *SD* (dashed lines) or quadratic form testing, abbreviated as *Q* (dotted lines), is defined analogously.

**Figure 15:**
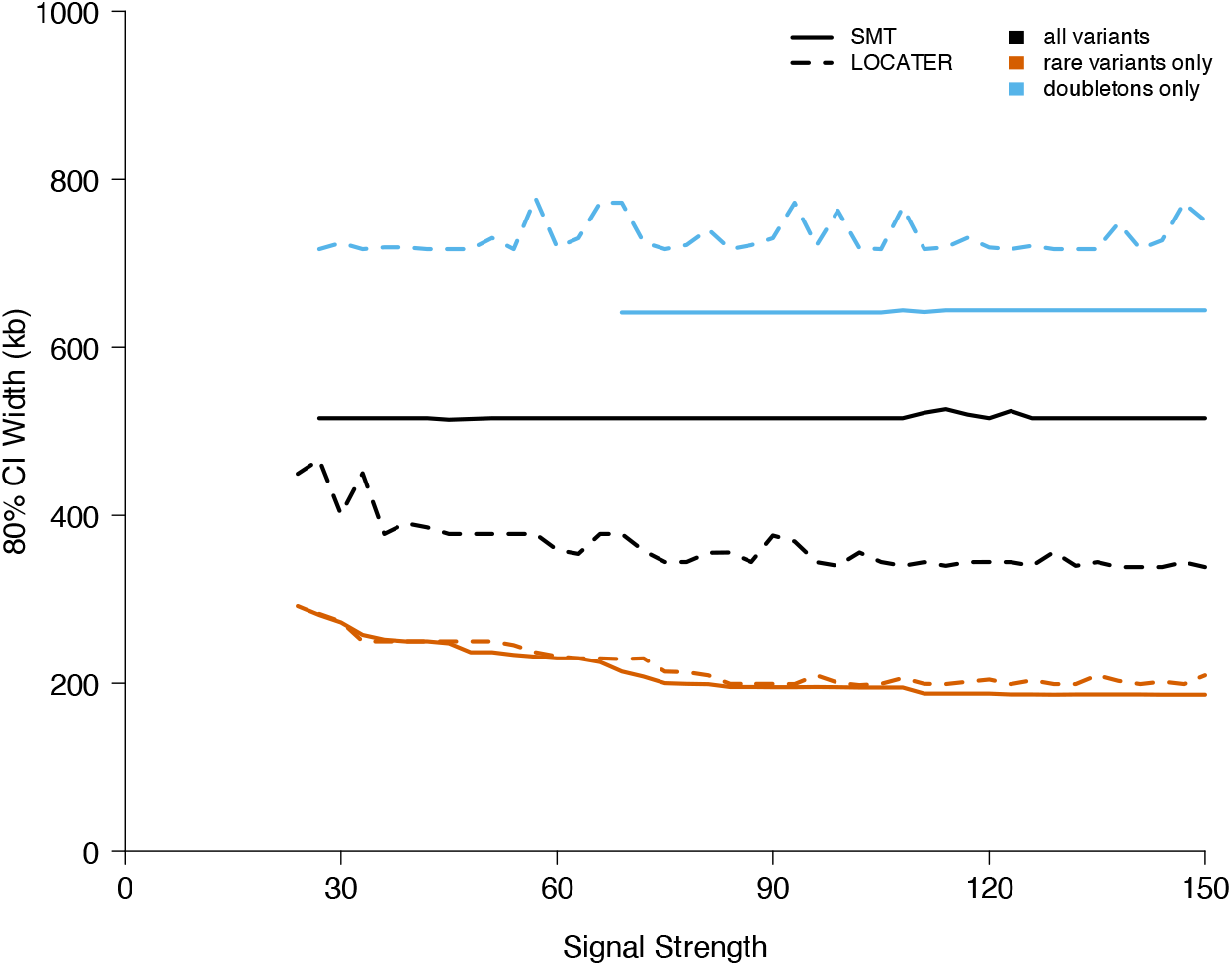
Width of confidence intervals (CI), centered on the position with the largest signal, that cover the midpoint of the causal region in 80% of simulations where there are 3 causal variants, all hidden. In simulations where multiple variants are tied to have the largest association signal, we take the distance to the midpoint of the causal region to be the average distance from each of the tied variants.

**Figure 16:**
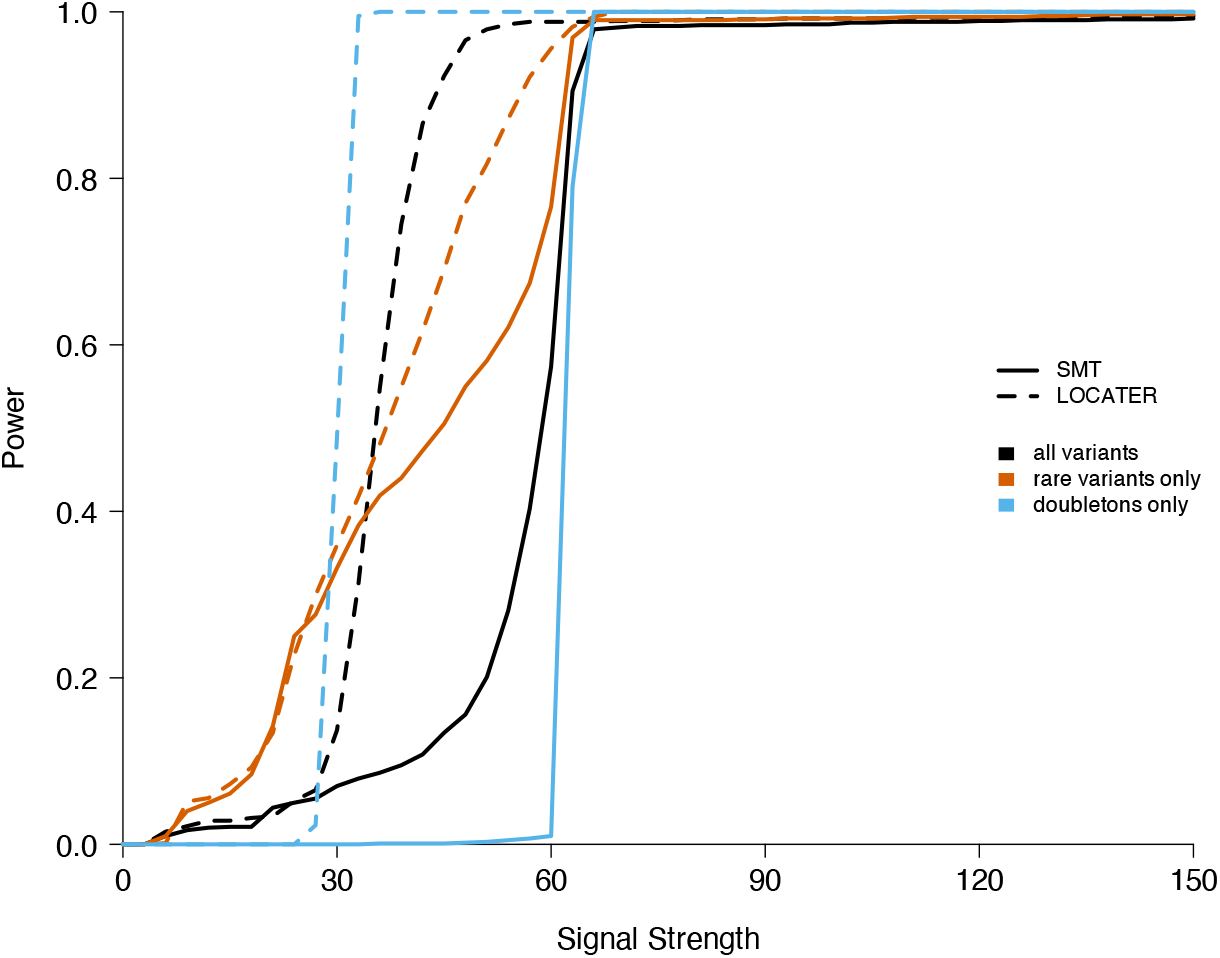
Power comparison between SMT and LOCATER in simulations where there are 9 causal variants, all observed.

**Figure 17:**
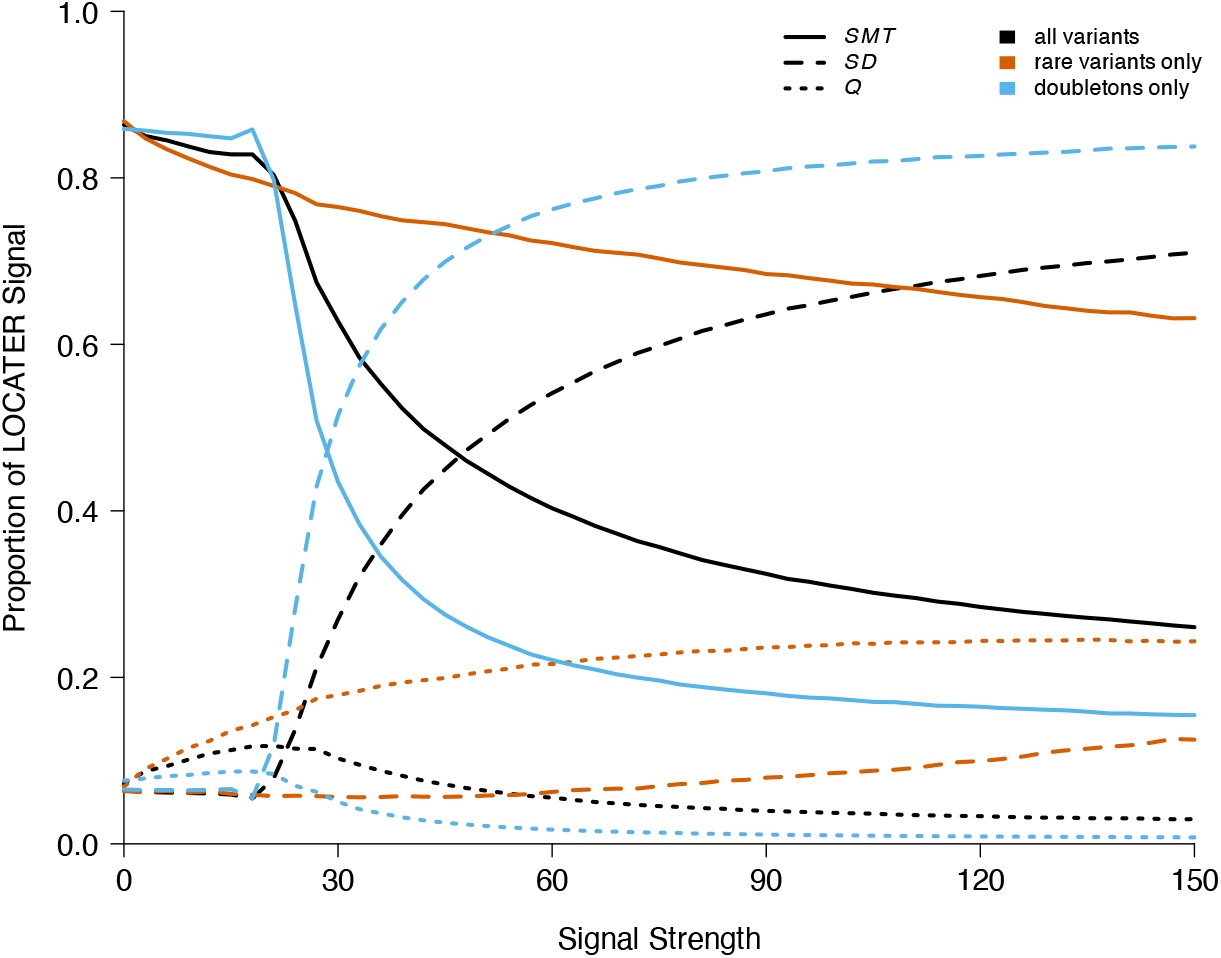
Average proportion of signal driven by each sub-test within LOCATER in simulations where there are 9 causal variants, all observed. For every simulation, the proportion of signal attributed to the *SMT* part of LOCATER (solid lines) defined as 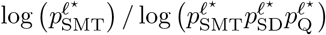 where *ℓ*^⋆^ is the locus *ℓ* that achieves the largest combined LOCATER p-value. The proportion of signal attributed to *SD* (dashed lines) or quadratic form testing, abbreviated as *Q* (dotted lines), is defined analogously.

**Figure 18:**
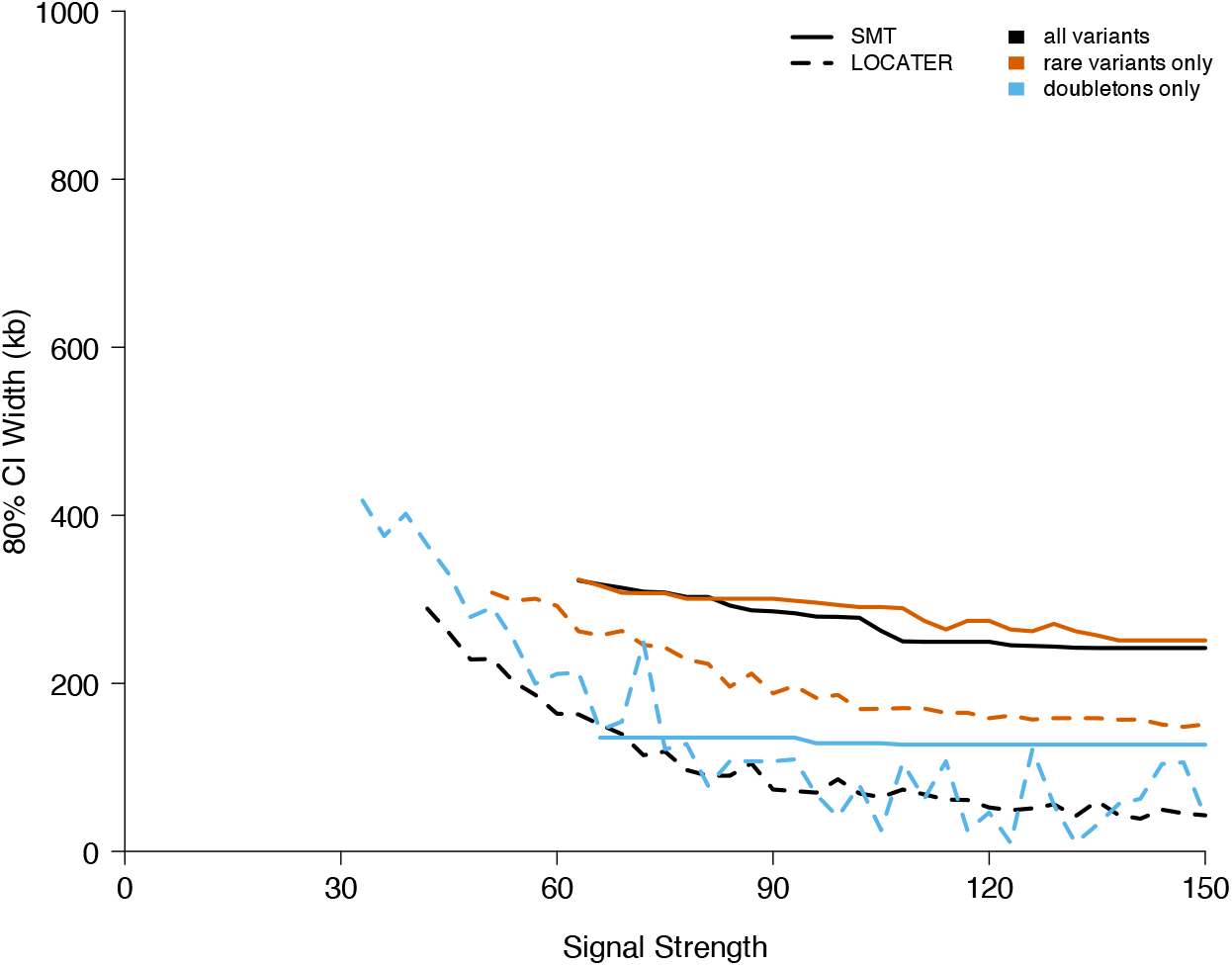
Width of confidence intervals (CI), centered on the position with the largest signal, that cover the midpoint of the causal region in 80% of simulations where there are 9 causal variants, all observed. In simulations where multiple variants are tied to have the largest association signal, we take the distance to the midpoint of the causal region to be the average distance from each of the tied variants.

**Figure 19:**
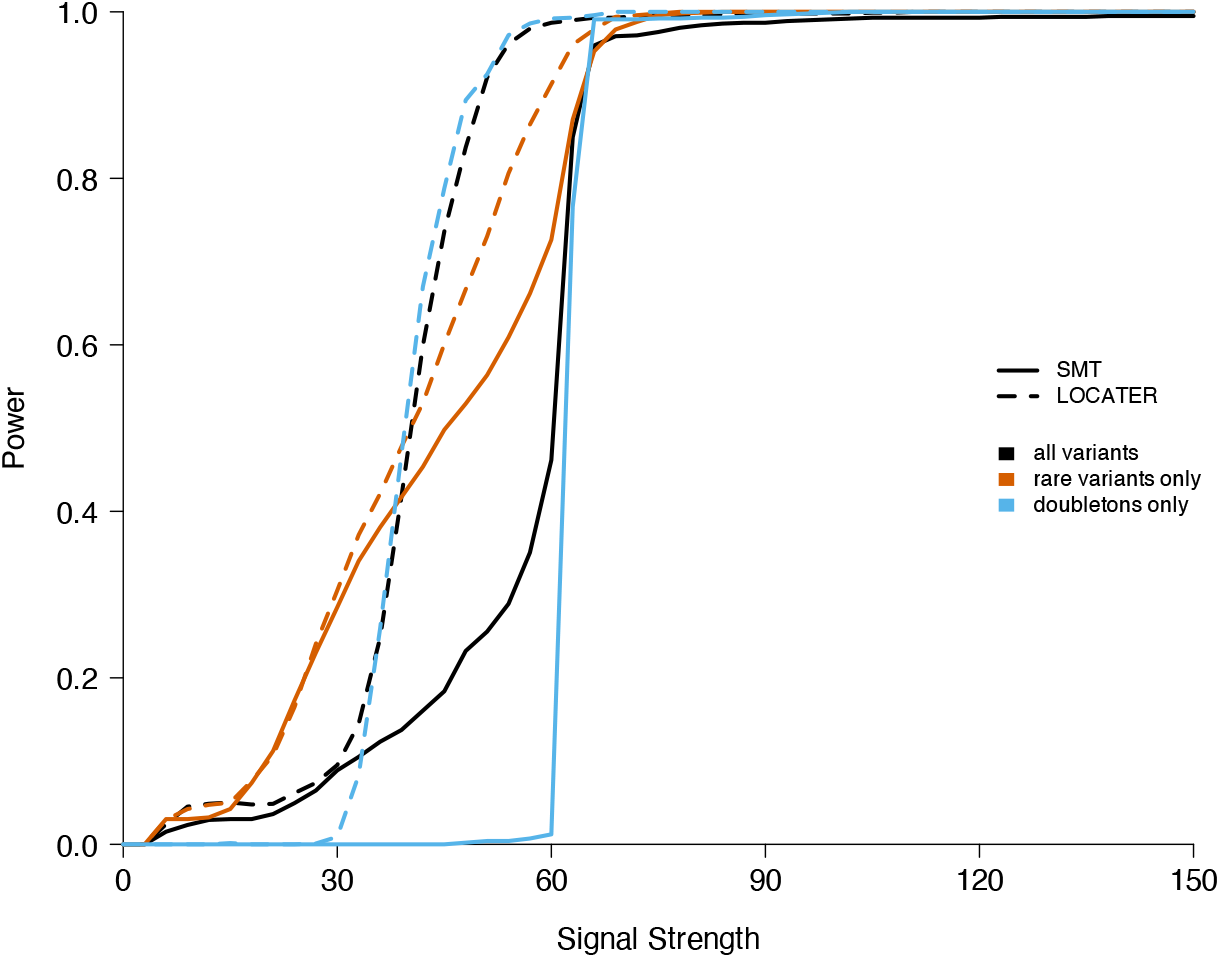
Power comparison between SMT and LOCATER in simulations where there are 9 causal variants, all hidden.

**Figure 20:**
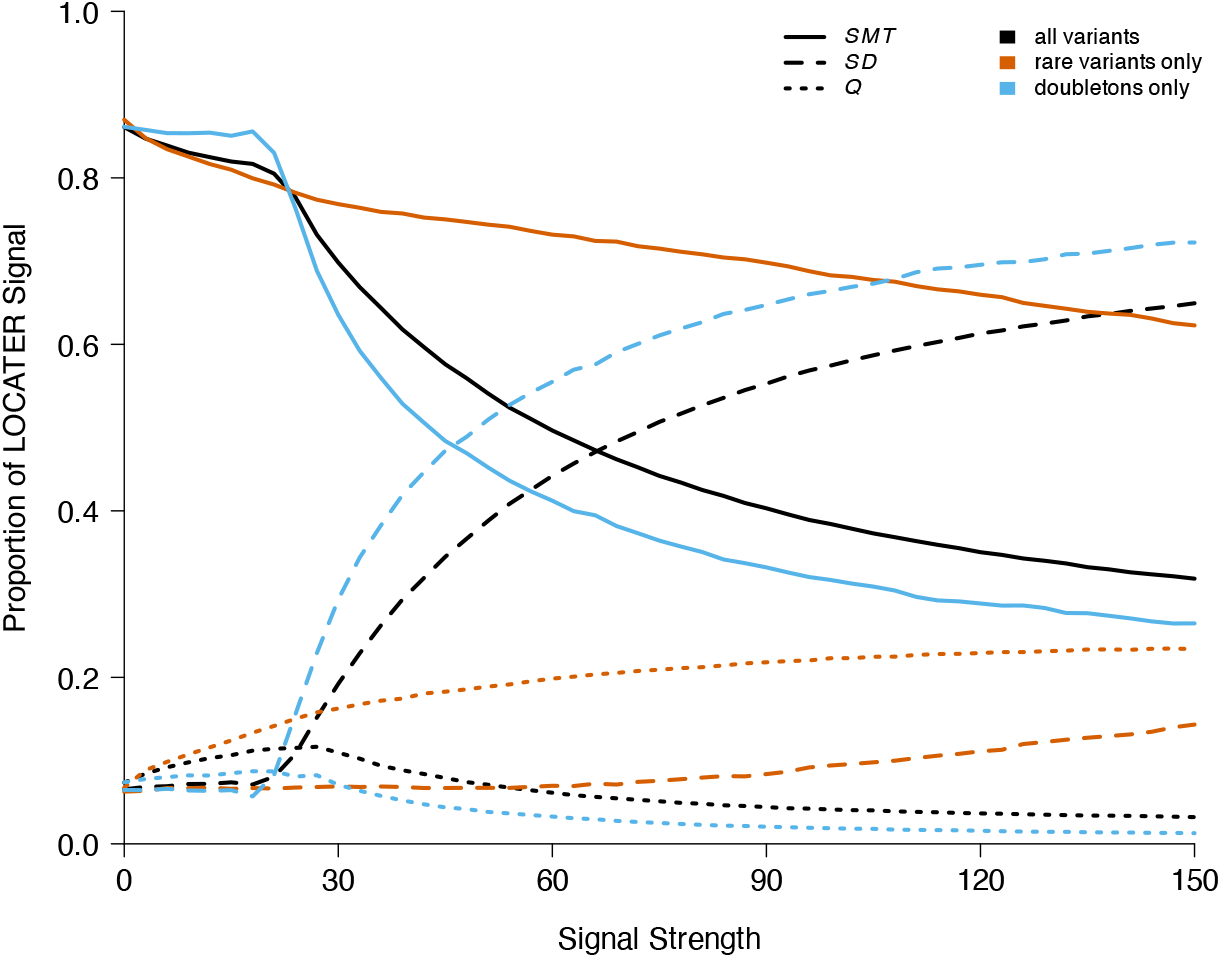
Average proportion of signal driven by each sub-test within LOCATER in simulations where there are 9 causal variants, all hidden. For every simulation, the proportion of signal attributed to the *SMT* part of LOCATER (solid lines) defined as 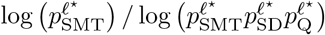 where *ℓ*^⋆^ is the locus *ℓ* that achieves the largest combined LOCATER p-value. The proportion of signal attributed to *SD* (dashed lines) or quadratic form testing, abbreviated as *Q* (dotted lines), is defined analogously.

**Figure 21:**
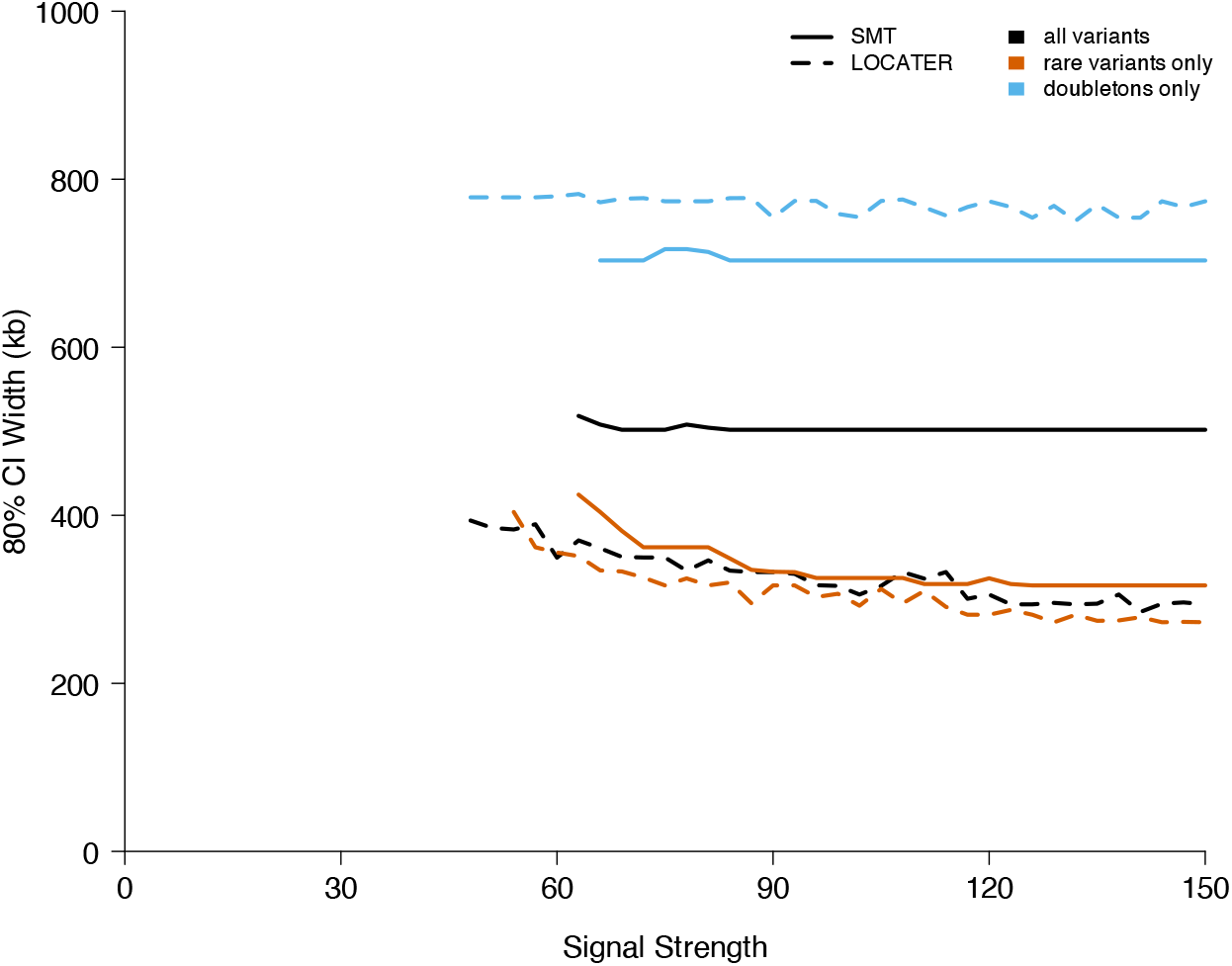
Width of confidence intervals (CI), centered on the position with the largest signal, that cover the midpoint of the causal region in 80% of simulations where there are 9 causal variants, all hidden. In simulations where multiple variants are tied to have the largest association signal, we take the distance to the midpoint of the causal region to be the average distance from each of the tied variants.

**Figure 22:**
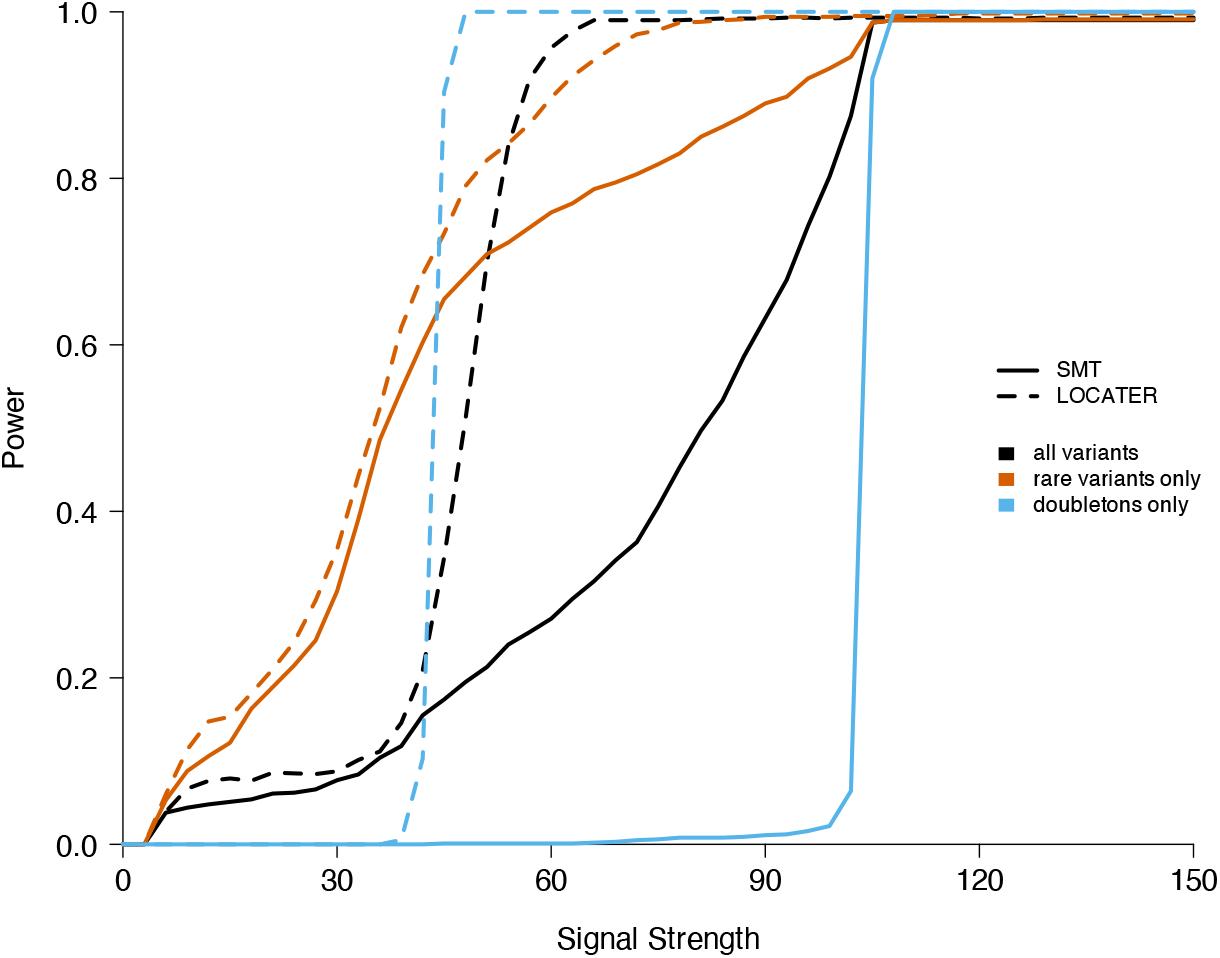
Power comparison between SMT and LOCATER in simulations where there are 15 causal variants, all observed.

**Figure 23:**
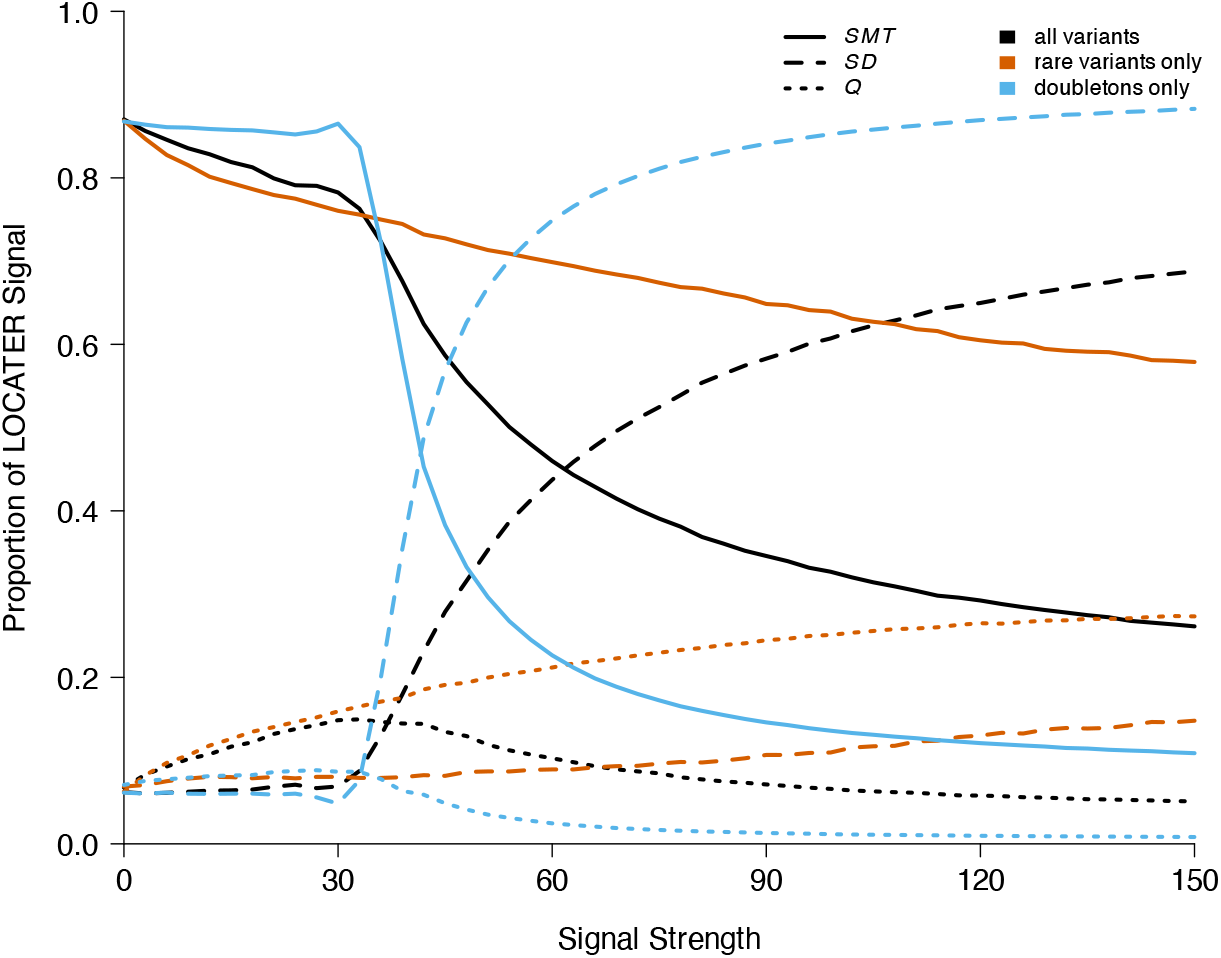
Average proportion of signal driven by each sub-test within LOCATER in simulations where there are 15 causal variants, all observed. For every simulation, the proportion of signal attributed to the *SMT* part of LOCATER (solid lines) defined as 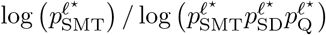 where *ℓ*^⋆^ is the locus *ℓ* that achieves the largest combined LOCATER p-value. The proportion of signal attributed to *SD* (dashed lines) or quadratic form testing, abbreviated as *Q* (dotted lines), is defined analogously.

**Figure 24:**
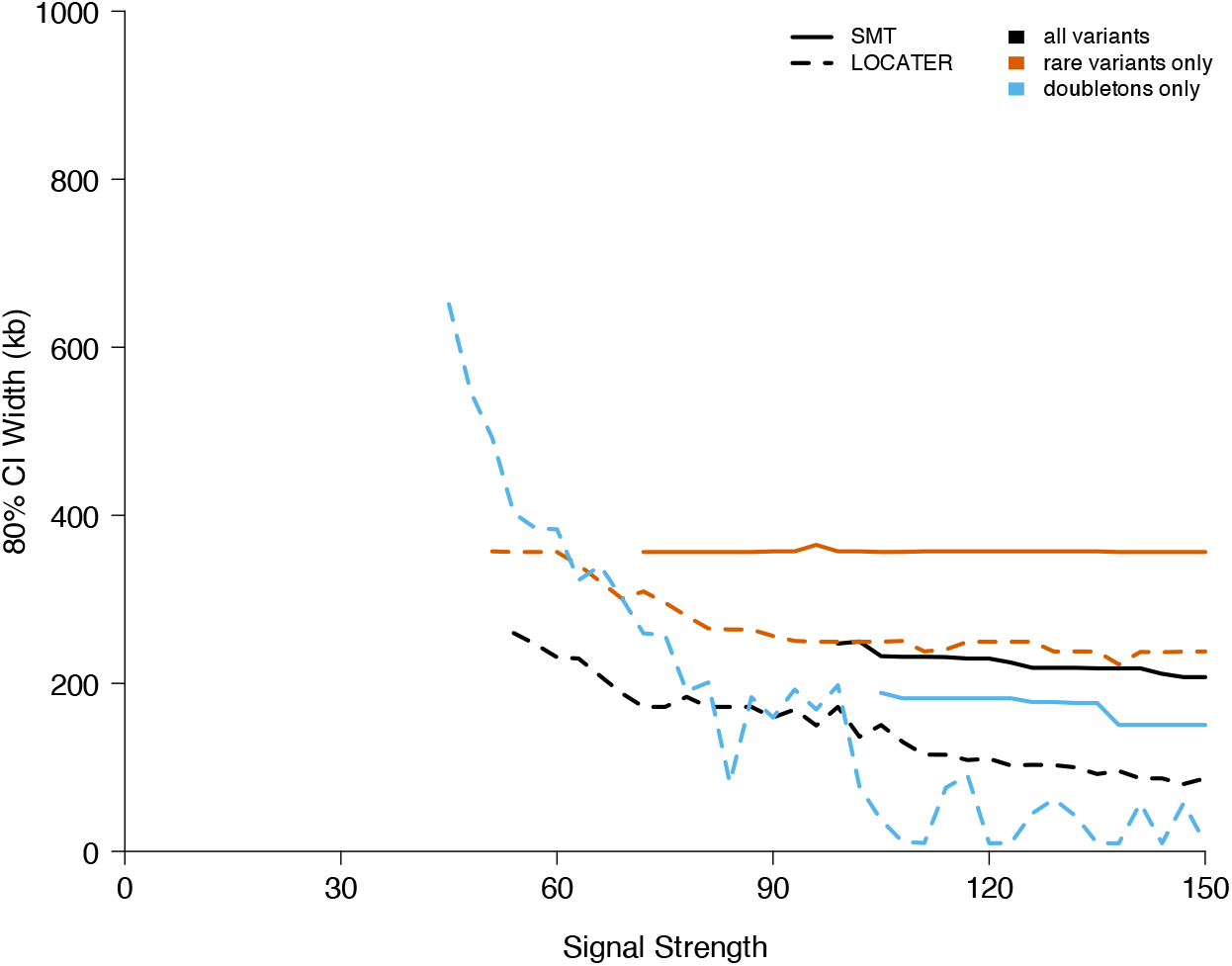
Width of confidence intervals (CI), centered on the position with the largest signal, that cover the midpoint of the causal region in 80% of simulations where there are 15 causal variants, all observed. In simulations where multiple variants are tied to have the largest association signal, we take the distance to the midpoint of the causal region to be the average distance from each of the tied variants.

**Figure 25:**
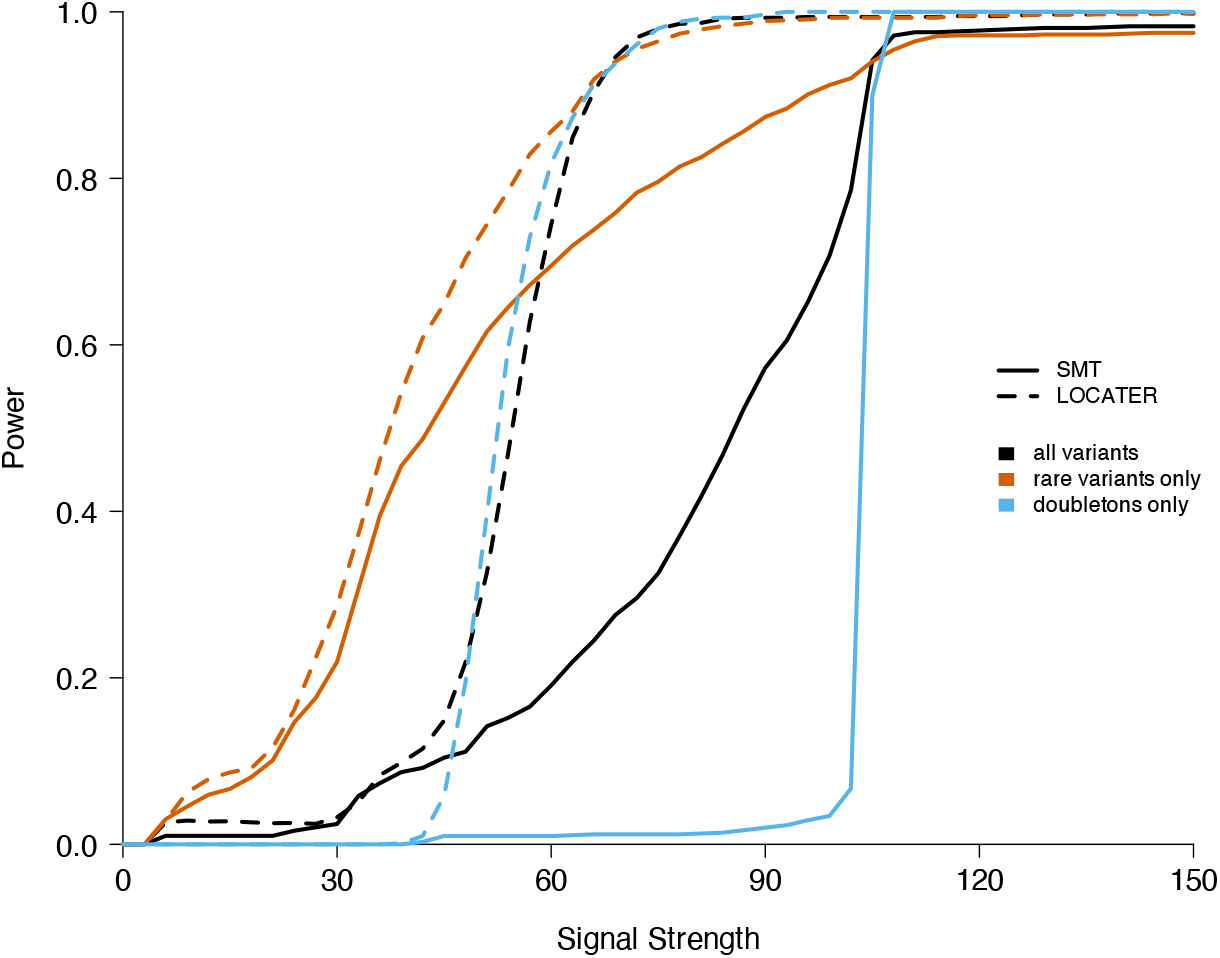
Power comparison between SMT and LOCATER in simulations where there are 15 causal variants, all hidden.

**Figure 26:**
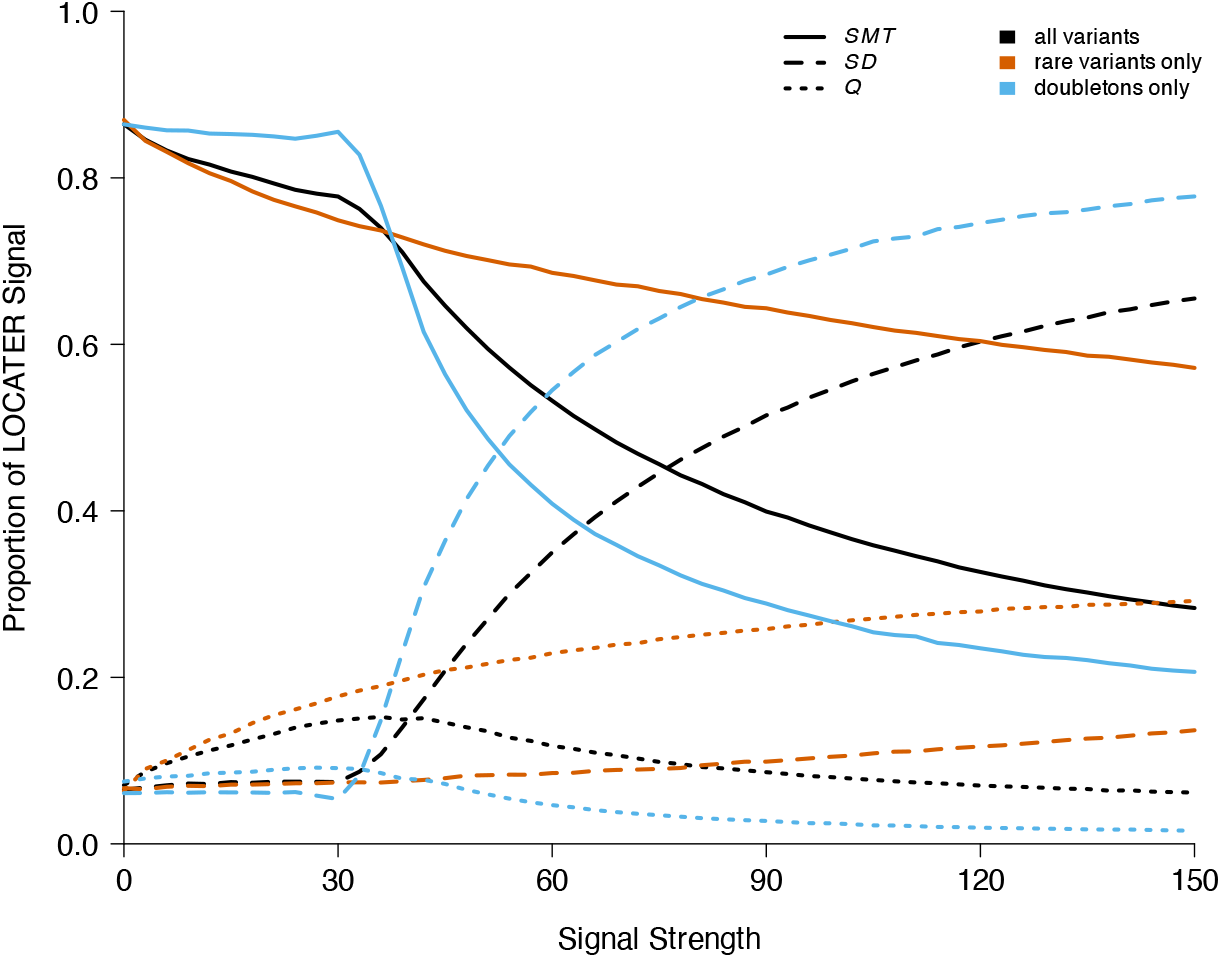
Average proportion of signal driven by each sub-test within LOCATER in simulations where there are 15 causal variants, all hidden. For every simulation, the proportion of signal attributed to the *SMT* part of LOCATER (solid lines) defined as 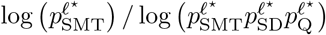 where *ℓ*^⋆^ is the locus *ℓ* that achieves the largest combined LOCATER p-value. The proportion of signal attributed to *SD* (dashed lines) or quadratic form testing, abbreviated as *Q* (dotted lines), is defined analogously.

**Figure 27:**
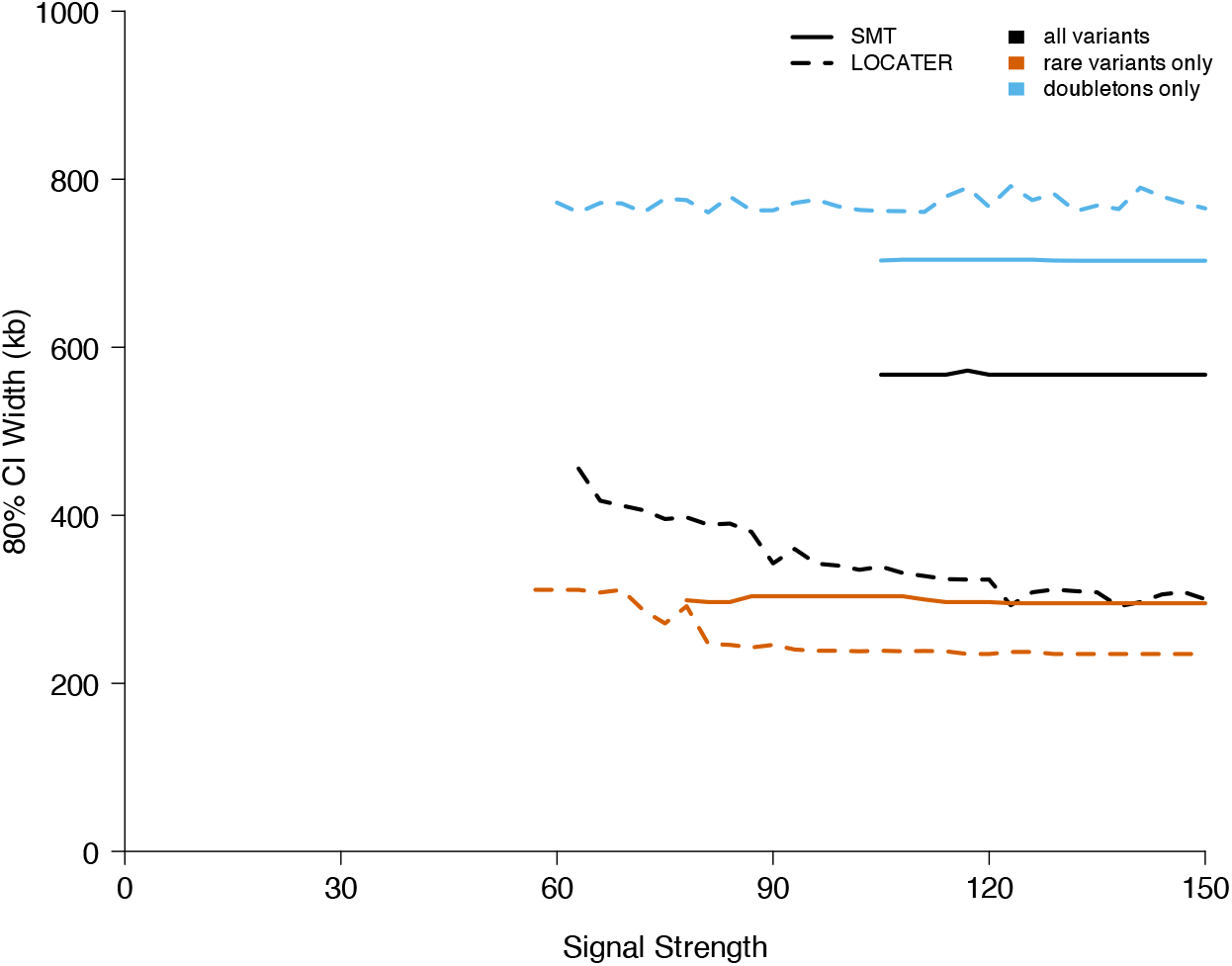
Width of confidence intervals (CI), centered on the position with the largest signal, that cover the midpoint of the causal region in 80% of simulations where there are 15 causal variants, all hidden. In simulations where multiple variants are tied to have the largest association signal, we take the distance to the midpoint of the causal region to be the average distance from each of the tied variants.

### 5.4 Power Curves: 100 kb Causal Window

**Figure 28:**
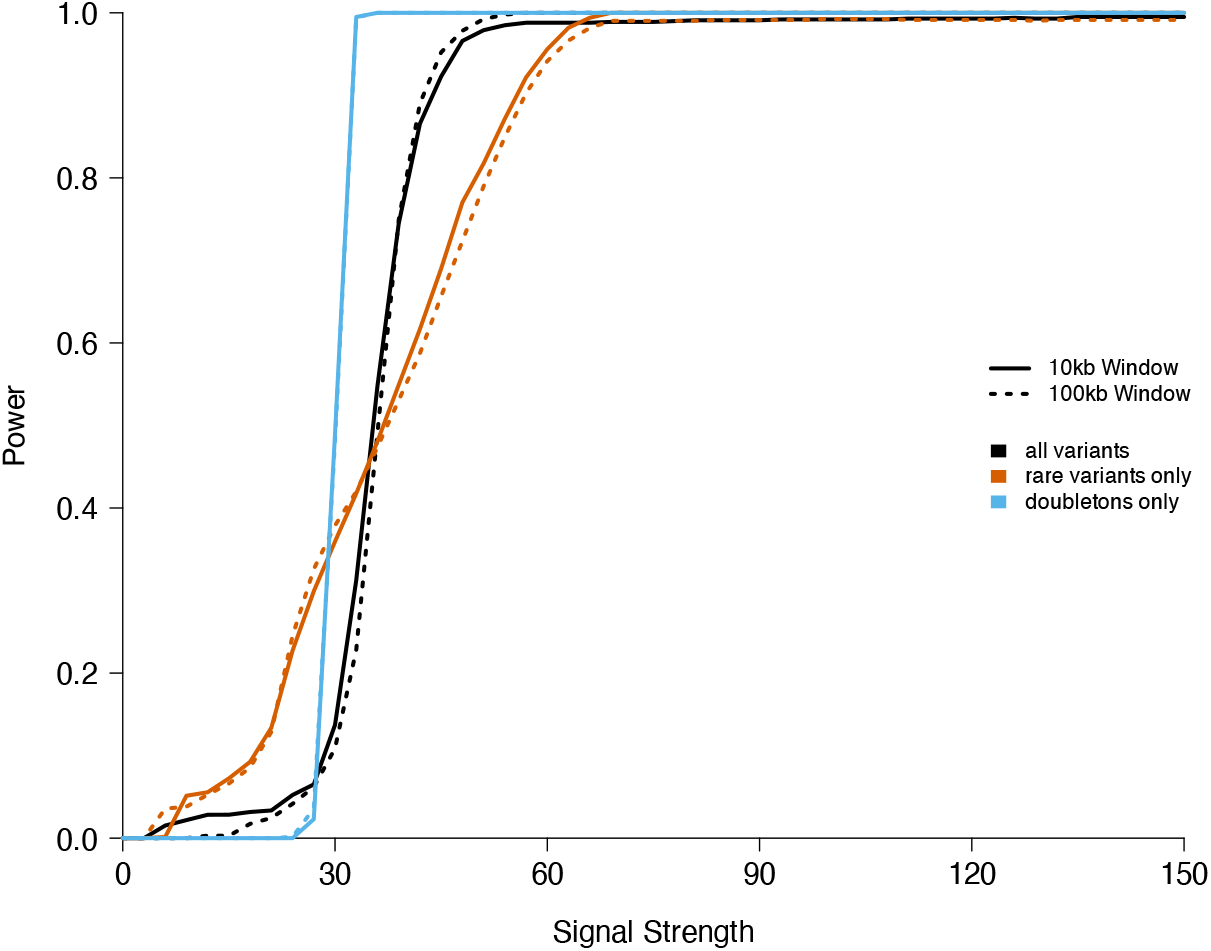
Power comparison between LOCATER when all causal variants are within a 10 kb causal region (solid lines) and LOCATER when all causal variants are within a 100 kb causal region (dotted lines) in simulations where there are 9 causal variants, all observed.

**Figure 29:**
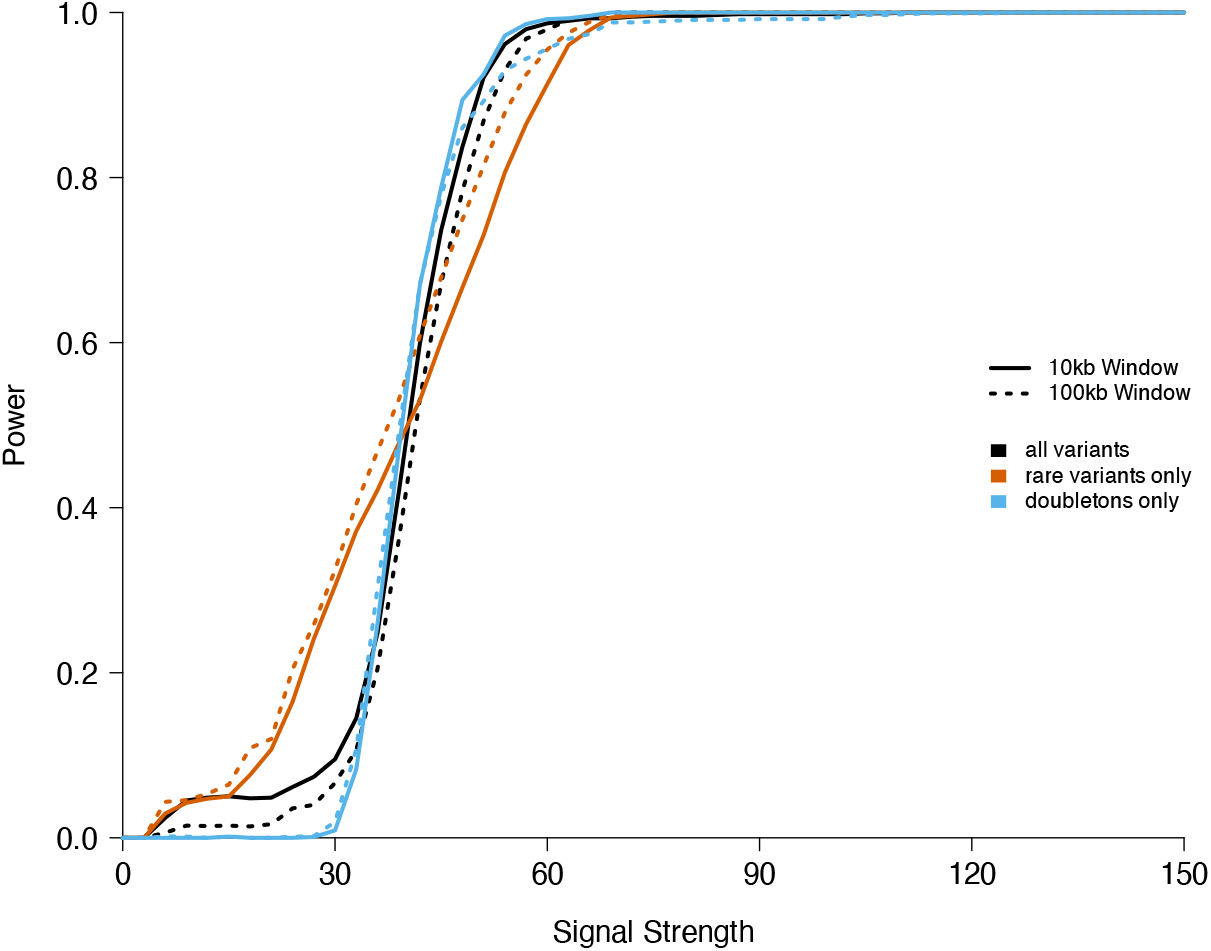
Power comparison between LOCATER when all causal variants are within a 10 kb causal region (solid lines) and LOCATER when all causal variants are within a 100 kb causal region (dotted lines) in simulations where there are 9 causal variants, all hidden.

## 6 Supplementary Methods

### 6.1 Running kalis

As described in [29, 13, 15], the LS model requires a pre-defined recombination map and two model parameters: *μ* governing the probability of mutations and *N*_*e*_ recombination events. Throughout this paper, we provided `kalis` with the true recombination map under which haplotypes were simulated. Throughout we set *μ* = 10^−4^ and *N*_*e*_ = 10^−16^.

### 6.2 Connecting our Generalized eGRM to the standard eGRM

Here we explain the connection between the generalized eGRM matrix defined in Equation (10) to the standard eGRM, as presented in Equation 1 in [17]. For clarity, throughout this section we will assume that we are considering a particular target locus *ℓ* and suppress indexing objects by *ℓ*.

Let *n* be the number of samples in a genetic dataset. Let

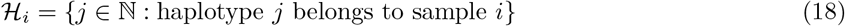

and ∆_*i*_ = |ℋ_*i*_| be the ploidy of sample *i* so that *N* = ∑_*i*_ ∆_*i*_ is the total number of haplotypes in the sample. Let *H* ∈ {0, 1}^*N*×*p*^ be the haplotype matrix encoding the carriers of *p* in a genomic region centered on locus *ℓ* or on an inferred local ancestral tree at *ℓ* as in [17]. Let 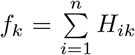 be the allele frequency of the *k*^*th*^ variant. Given a set of weights 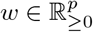 which may be based on inferred clade probabilities or otherwise, we start with a weighted haplotype relatedness matrix

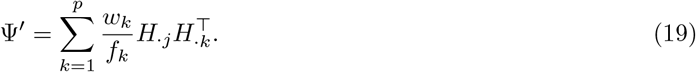

We can write this more compactly as 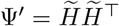 where 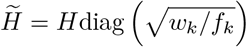. Assuming an additive model, we can collapse this weighted haplotype relatedness matrix to a weighted genotype relatedness matrix

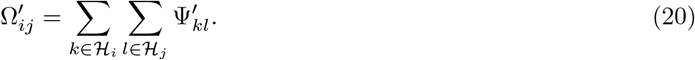

For simplicity, from here forward we will assume that (1) the ploidy of each sample ∆_*i*_ is the same constant ploidy ∆ for all samples *i* and (2) the rows and columns of Ψ^*′*^ are permuted such that haplotypes from the same sample are grouped together. This allows us to more succinctly write 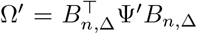 where *B*_*n*,∆_ = *I*_*n*×*n*_ ⊗ **1**_∆_ and ⊗ is the Kronecker product. This gives us

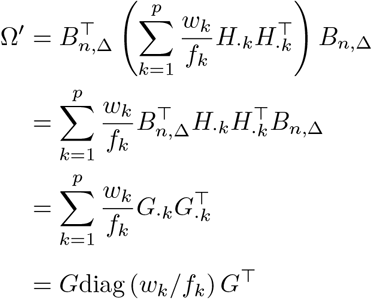

where *G* {0, 1, 2}^*n*×*p*^ is the genotype matrix corresponding to the *p* variants on the local tree. Now if we consider a model without any background covariates besides an intercept, we have 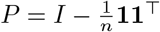.

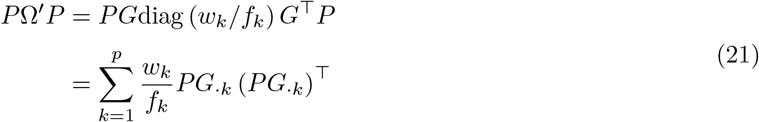

The standard GRM assumes that each variant is in Hardy–Weinberg equilibrium. So in order to compare Equation (21) to the standard GRM, let us also assume that each of our *p* variants is in perfect Hardy– Weinberg equilibrium. That means each centered genotype vector *PG*_·*k*_ = *G*_·*k*_ − 2*f*_*k*_**1**, allowing us to write

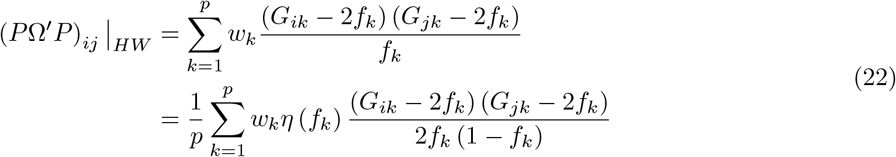

where *η* (*f*_*k*_) = 2*p* (1 − *f*_*k*_). The standard GRM defined in Equation 1 of [17] is just a simple special case of Equation (22) where *η* (*f*_*k*_) = 1 for all arguments *f*_*k*_ and *w*_*k*_ = 1 for all variants. The eGRM allows for non-equal weights *w*_*k*_. Some simple algebra shows that our choice of a 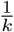 in our definition of *ψ* in Equation 9 corresponds to *η* (*f*_*k*_) = 2*p* (1 − *f*_*k*_). We chose this weighting in order to upweight variants that have low derived allele frequency. Under mild selection pressure, we expect very common derived alleles to have smaller phenotypic effects. With very minimal modification of our current implementation, any weighting function of the derived allele frequency *η* may be used. We plan to make such a general weighting function available in future versions of LOCATER.

Having rewritten the eGRM in terms of Ψ^*′*^, let us connect Ψ^*′*^ to Ψ from Equation (9), which we use in our generalized eGRM construction. Consider the special case where we have a chromosome without recombination so that our genetic distance matrix *d*^(*ℓ*)^ reflects a single marginal tree, making all clade calls are aligned. If we also set each *g*_*k*_ to the identity function and assume consistency across columns of our distance matrix so that 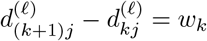 for all haplotypes *j* carrying a variant of frequency *k*, then Ψ as defined in Equation (9) would be symmetric and proportional to the the weighted haplotype matrix Ψ^*′*^ as defined in Equation (19). As a consequence, the similarity matrix we would obtain 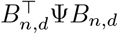 would equal 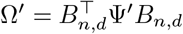. However, due to recombination, the distance matrix Ψ is not symmetric and the distances across columns may not be consistent with a single tree structure. Rather than attempting to synchronize the distances by clustering them into a tree structure, we trust the LS model, taking the distances as representing a weighted convolution of nearby tree structures.

### 6.3 Derivation of Robust Tail Approximation for Quadratic Forms: Accounting for Unmeasured Confounders

Consider the presence of some unobserved variable *B* ∈ ℝ^*n*^. Without loss of generality, assume 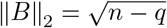 and *PB* = *B*. For some scalar *γ*, we have *Y* = *Aα*+*γB*+*σϵ*. Then our estimator 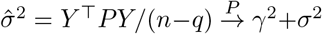. This gives us the null distribution

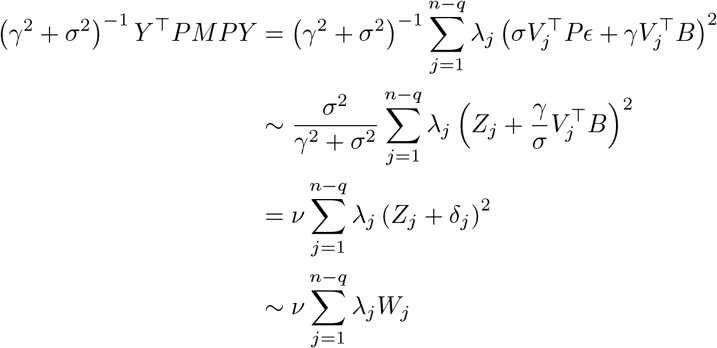

where 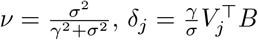, and each 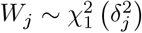, independent. The parameter *ν* ∈ [0, 1] captures the proportion of the variance of *Y* that is not explained by the unobserved variable *B* and each 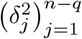 captures how correlated the clade structure encoded in *M* is with the unobserved variable *B*.

A typical clade matrix typically decays over the first ten or so eigenvalues and levels off to a plateau of eigenvalues that are all roughly of the same magnitude. The initial eigenvalues also tend to explain a small proportion of the variance of the matrix. This behavior is expected given the relative abundance of rare variants (small clades) which correspond to small eigenvalues. As a consequence, the contribution of *R*_*k*_ to the overall variance of the test statistic is much larger than the contribution of *T*_*k*_. The role of *T*_*k*_ is primarily to define the rate of decay in the tail. Let *k*^⋆^ denote the transition point in the spectrum to a plateau, defined as follows

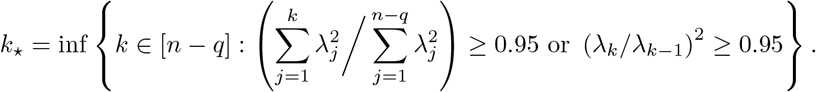

In order to ensure reliable estimation of the null distribution, assume that we compute the top *k* ≥ *k*^⋆^ eigenvalues of *PMP*, so we have

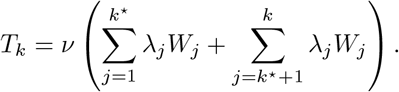

We can gather information about *ν* by looking at the distribution of this quadratic form test statistic at a grid of loci spaced far apart along the genome. Unfortunately, we cannot attempt to estimate each individual *δ*_*j*_ in a similar way since each specific *δ*_*j*_ depends on the precise *j*^*th*^ eigenvector *V*_*j*_ calculated at a given position. However, the symmetry of the multivariate normal distribution helps us tackle this problem.

If all of the eigenvalues, *λ*_*j*_ were equal, our null distribution would have a scaled non-central chi-square distribution with *n*−*q* degrees of freedom and a single non-centrality parameter 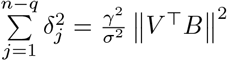. This collapse into a single non-centrality parameter means that we would not need to model each individual 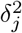, but rather just the average 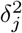. Since we expect that the typical angle between *B* and each *V*_*j*_ may be different for *V*_*j*_s capturing common variant structure versus rare variant structure, we replace each 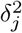 such that

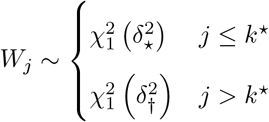

This leaves us with a null distribution with three scalar inflation parameters, 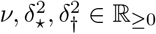.

We can use tr (*PMP*) and 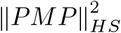 to calculate the mean and variance of *R*_*k*_. Using those moments, we approximate *R*_*k*_ with a shifted difference of chi-square random variables. Let

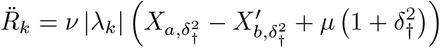

where 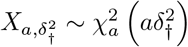 is distributed chi-squared with *a* degrees of freedom and non-centrality parameter 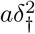 and independent of 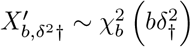.

We will select parameters *a, b, μ* so that 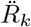 matches *R*_*k*_ on the first two moments for all values of *ν* and *δ*_†_. We can do this using matrix traces to exactly calculate 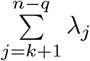 and 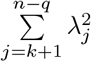. If we then set the first moments equal we get

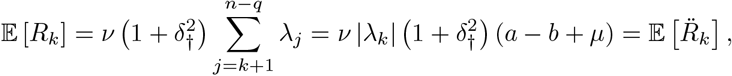

yielding the constraint

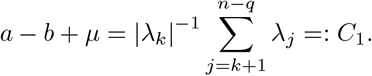

Similarly, setting the variances equal we have

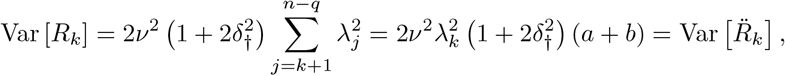

which yields the constraint,

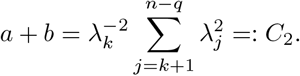

With these constraints, we select *a, b, μ* as follows

1. If *C*_1_ ∈ [−*C*_2_, *C*_2_], we set *a* = (*C*_2_ + *C*_1_) */*2 and then set *b* = *C*_2_ − *a* and *μ* = 0.
2. If *C*_1_ > *C*_2_, we set *a* = *C*_2_, *b* = 0 and *μ* = *C*_1_ − *C*_2_.
3. If *C*_1_ < −*C*_2_, we set *b* = *C*_2_, *a* = 0 and *μ* = *C*_1_ + *C*_2_.

This specification minimizes the contribution of *μ* to the mean relative to *a* and *b* while respecting our constraints. This yields a final null distribution

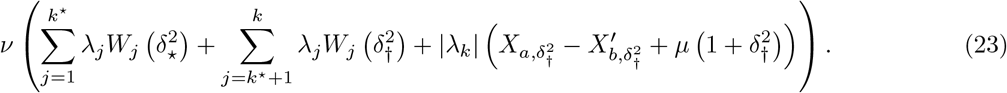

Critically, this parameterization of the null distribution is expressed in two sets of orthogonal parameters. The first set are our spectral parameters 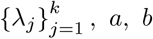, and *μ* corresponding to the spectrum of *PMP*. The second are our mis-specification parameters 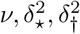 corresponding to features of the confounding process. This means that, given the phenotype vector *Y* and the spectral parameters at a given locus, we can readily recompute the observed p-value for that locus for any set of misspecification parameters. Thus, if we see inflation in the Q-Q plot of observed quadratic form p-values for a given phenotype, we can readily adjust the mis-specification parameters and recalculate our p-values in order to correct and calibrate the Q-Q plot.

### 6.4 Fast Trace Calculation

Here we present our trace calculation approach for a *symmetric* matrix *M* ∈ ℝ^*n*×*n*^. Let *B* = *I*_*n*_ − *QZ* where *Q* ∈ ℝ^*n*×*m*^ and *Z* ∈ ℝ^*m*×*n*^. Our goal is to calculate 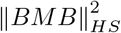 and diag (*BMB*) efficiently using a set of worker nodes. First we compute *J* = *ZM* ∈ ℝ^*m*×*n*^ (*mn*^2^ FLOPs). Second we compute *X* = *Q*(*JQ*) − *MQ* ∈ ℝ^*n*×*m*^ (*mn*^2^ + 2*nm*^2^ + *nm* FLOPs). Then map out the relevant sub-blocks of *Q, Z, J, X* to each worker as follows:

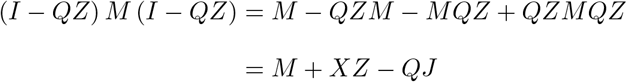

So,

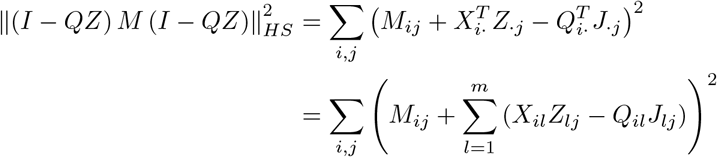

This accumulation requires a total of (4 + 1)^2^ FLOPs. Since diag 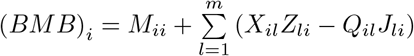, diag (*BMB*) is easily obtained as a by-product of this calculation. Taking the sum of those diagonal elements to obtain tr (*BMB*) requires *n* − 1 FLOPs. Adding the FLOPs together, we obtain a total of (6*m* + 1)*n*^2^ + 2*nm*^2^ + *nm* + *n* − 1 FLOPs. Since *m* ≥ 1, this is upper bounded by 7*mn*^2^ + 3*nm*^2^. Connecting this result back to our notation in Section 6.3, this means we can calculate our desired traces *η*_1_ and *η*_2_ with fewer than 7(*q* + 1)*n*^2^ + 3*n*(*q* + 1)^2^ FLOPs where *q* is the number of columns in our background covariate matrix *A*.

If *M* is distributed in columns across nodes, we can accelerate the above calculation by exporting *W* =[*X* −*Q]* ∈ ℝ^*n*×2*m*^ to every node and the appropriate columns of 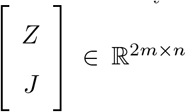 to their corresponding nodes.

### 6.5 Haplotype Simulation

We simulated haplotypes from three 1000 Genomes Populations – Yoruba, Han Chinese, and Central European – using a demographic model adapted from the `msprime` [36] demography tutorial [38], which itself was based on the population parameters related those three populations presented in [39]. We only made one modification to the demographic model specified in [38]: we use the Discrete Time Wright–Fisher model for the first 100 generations into the past before reverting back to the classic Hudson model as proposed in [40]. The mutation rate was set to 1.2×10^−8^.

Each 1 Mb region of simulated haplotypes was generated using a population-specific human recombination map estimated by pyrho [41]. Explicitly, in each simulation, we randomly selected one of our three 1000 Genomes populations (YRI, CHB, or CEU) and 1 Mb segment from genome (excluding heterochromatic regions), then we loaded the recombination map estimated for that population and region by `pyrho`.

### 6.6 Phenotype Rank Matching

Before we run LOCATER, as has become standard practice in testing quantitative traits, we require that the phenotype residuals be rank normalized after fitting background covariates. The theory underlying SD requires that the phenotypes be independent under 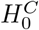. Standard inverse-rank-normalization maps phenotype residuals to a fixed grid of values based on the Gaussian CDF, which in turn induces dependency between phenotypes. In order to avoid inducing that dependency, we substitute each phenotype residual with a rank-matched Gaussian random variable. This achieves a rank normalization that is very similar to inverse-rank-normalization but with a small amount of noise added in order to make the normalized phenotypes truly independent under the null hypothesis.

Explicitly, given some raw phenotype vector 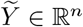 and a matrix of background covariates *A* ∈ ℝ^*n*×*q*^ (including an intercept), we begin by calculating the residuals 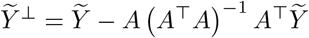 of the standard ordinary least squares model

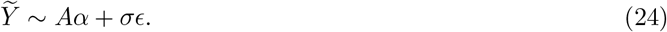

Then we find the permutation *π* : [*n*] → [*n*] that orders the residuals 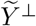 such that 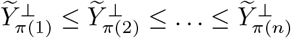. We simulate a new vector of Gaussian phenotypes, *Z* ∼ *N* (0, *I*_*n*_) and likewise find the permutation *π*^*′*^ : [*n*] → [*n*] that orders the entries of *Z* such that *Z*_*π ′*(1)_ ≤ *Z*_*π ′*(2)_ ≤ … ≤ *Z*_*π ′*(*n*)_. Then we obtain our normalized vector of rank-matched phenotypes, *Y* ∈ ℝ^*n*^, by assigning 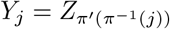.

